# Muscle specific stress fibers give rise to sarcomeres and are mechanistically distinct from stress fibers in non-muscle cells

**DOI:** 10.1101/235424

**Authors:** Aidan M. Feinx, Nilay Taneja, Abigail C. Neininger, Mike R. Visetsouk, Benjamin R. Nixon, Annabelle E. Manalo, Jason R. Becker, Scott W. Crawley, David M. Bader, Matthew J. Tyska, Jennifer H. Gutzman, Dylan T. Burnette

## Abstract

The sarcomere is the basic contractile unit within cardiomyocytes driving heart muscle contraction. We sought to test the mechanisms regulating thin (i.e., actin) and thick (i.e., myosin) filament assembly during sarcomere formation. Thus, we developed an assay using human cardiomyocytes to test *de novo* sarcomere assembly. Using this assay, we report a population of muscle-specific stress fibers are essential sarcomere precursors. We show sarcomeric actin filaments arise directly from these muscle stress fibers. This process requires formin-mediated but not Arp2/3-mediated actin polymerization and nonmuscle myosin IIB but not non-muscle myosin IIA. Furthermore, we show a short species of β cardiac myosin II filaments grows to form ~1.5 long filaments that then “stitch” together to form the stack of filaments at the core of the sarcomere (i.e., A-band). Interestingly, these are different from mechanisms that have previously been reported during stress fiber assembly in non-muscle cells. Thus, we provide a new model of cardiac sarcomere assembly based on distinct mechanisms of stress fiber regulation between non-muscle and muscle cells.

## Introduction

At its core, a sarcomere is composed of “thick” myosin II filaments, and “thin” actin filaments (Figure 1A) (Au, 2004). The proper establishment of cardiac sarcomeres during development and their subsequent maintenance are critical for heart function. Indeed, mutations in sarcomeric proteins, such as actin and myosin II, drive devastating disease states such as dilated and hypertrophic cardiomyopathies (Bonne et al., 1998; Ho, 2010; Hughes and McKenna, 2005; Watkins et al., 2011). A testable platform to investigate mechanisms of sarcomere assembly would illuminate how sarcomeres arise in both normal and disease states. Unfortunately, no such assay exists. Previous studies investigating sarcomere assembly have utilized a number of model systems, including primary isolated cardiomyocytes from chick embryos and neonatal rats. These systems have provided important descriptive ground-work to start elucidating the mechanisms of sarcomere assembly. However, use of primary cell culture to investigate sarcomere assembly has a number of serious experimental caveats that compromised mechanistic studies of this process. First, isolated cardiomyocytes already contain sarcomere structures, making it difficult to separate *de novo* sarcomere assembly from homeostasis (i.e., turnover) of existing sarcomeres (Iskratsch et al., 2010; Rhee et al., 1994; Taniguchi et al., 2009). Second, long term culture, and thus extended experiments have proven difficult as isolated cardiomyocytes tend to die and/or de-differentiate in long term culture (Zhang et al., 2010). Third, and most importantly, these systems are genetically complicated, making mechanistic and live-cell experimentation both technically difficult and limited. Despite these drawbacks, previous studies have made important descriptive discoveries, which have led to a number of models of sarcomere assembly (Sanger et al., 2005).

**Figure 1:**
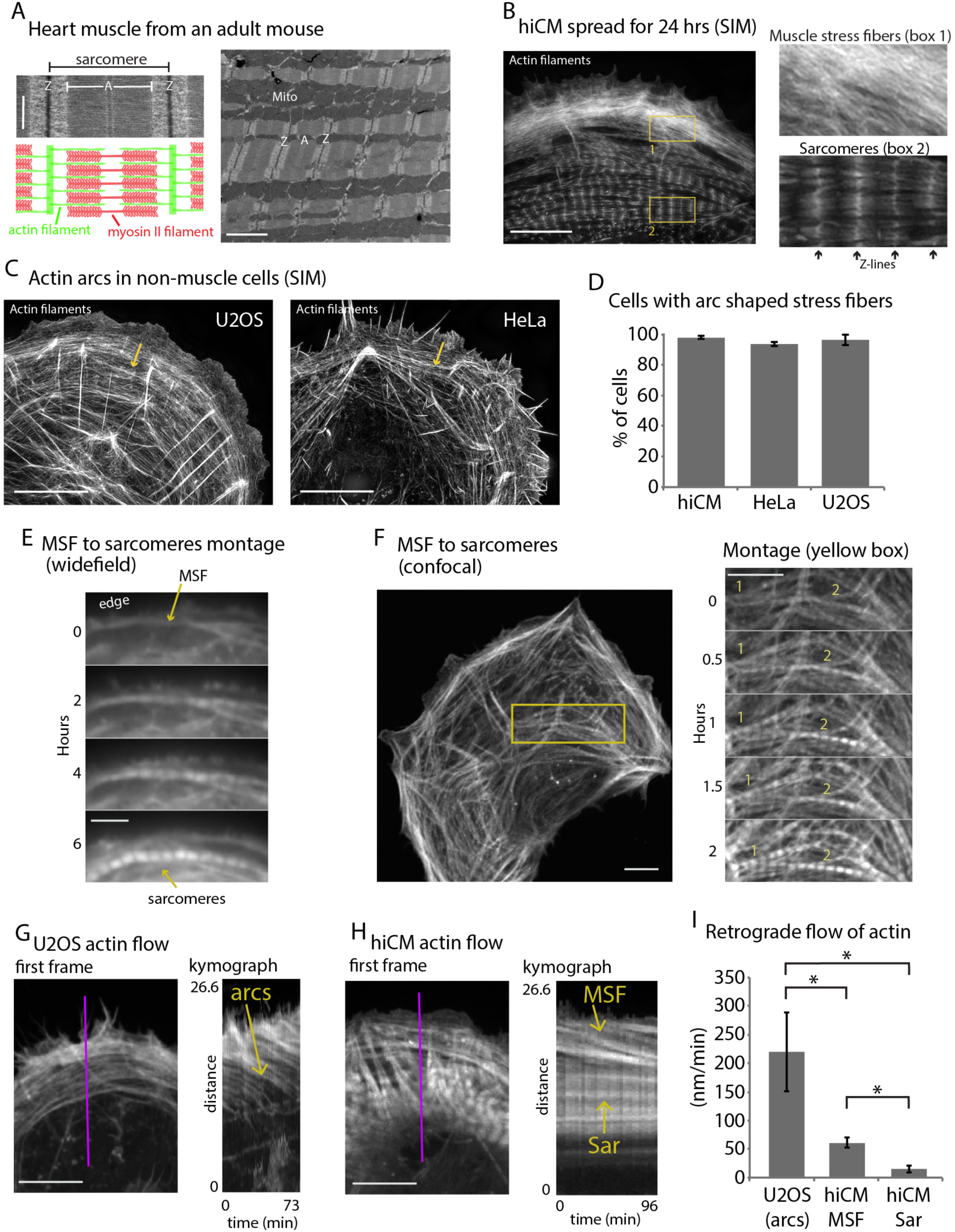
Sarcomeres arise directly from Muscle Stress Fiber (MSF) precursors. (A) Electron microscopy (EM) schematic of a cardiac sarcomere from adult mouse. Electron dense regions on border of sarcomere are Z-discs (Z), while the core of the sarcomere is composed of thin actin filaments and thick myosin II filaments (A). Multiple sarcomeres aligned adjacently form a myofibril (lower mag EM, right). (B) hiCM allowed to spread for 24 hours following plating. Notice the clear stress fiber and sarcomere-like actin organization at the front and rear of the cell in box 1 and 2, respectively. (C) Spread U2-OS cell (left) and HeLa cell (right) displaying prominent actin arc stress fibers behind leading edge of cell (yellow arrow). (D) Percentage of hiCMs, U2-OS, and HeLa cells with actin arc stress fibers. hiCMs; 1372 cells over 3 experiments. U2OS; 37 cells over 4 experiments. HeLa; 186 cells over 4 experiments. (E) Wide-field time lapse of hiCM transfected with Lifeact-mEmerald to visualize actin. MSF at front of hiCM undergoes retrograde flow and acquires sarcomeres (arrow). (F) Laser-scanning confocal microscopy of MSF to sarcomere transition. High magnification montage on right from blue box in low mag still image (left) (G) Still of U2-OS cell expressing Lifeact-mEmerald (left). Kymograph (right) taken from purple line of left image. Note robust movement of actin arc stress fibers (yellow arrow). (H) Still of hiCM expressing Lifeact-mEmerald (left). Kymograph (right) taken from purple line of left image. Note slower movement of MSF in hiCM compared to actin arcs in U2-OS cell (Fig. 1G), and stationary nature of sarcomeres. (I) Quantification of actin stress fiber translocations rates in U2-OS cells and hiCMs. U2OS; 5 cells over 4 experiments. hiCMs; 12 cells over 3 experiments. Scale Bars; (A) 500 nm high mag, 2 μm low mag (B), (C), (E) middle and right, (F), (G) 10 μm, (E) left 5 μm.

Studies from explanted chick cardiomyocytes have described the molecular motor, non-muscle myosin IIB (NMIIB), and the actin filament bundling protein, α-actinin 2, containing actin stress fibers located parallel to the edge of spreading cardiomyocytes, with organized sarcomere structures in the cell body (Rhee et al., 1994). Non-muscle myosin IIA (NMIIA) was reported to not localize to these stress fibers (Sanger et al., 2005). These studies proposed that the NMIIB stress fibers, referred to as “pre-myofibrils”, found at the edge gave rise to the sarcomere structures towards the cell body (Rhee et al., 1994; Sanger et al., 2005). These studies further show pre-myofibrils are largely devoid of sarcomere specific proteins (such as titin), which the sarcomere structures towards the cell body contain (Du et al., 2008; Rhee et al., 1994). Though a pre-myofibril structure containing actin, NMIIB, *α*-actinin has never been directly shown to transition into a sarcomere containing myofibril, previous live cell data using fluorescently tagged injected *α*-actinin has shown “Z-bodies” at the edge come together as the Z-bodies undergo retrograde flow and assemble into existing sarcomeres (Dabiri et al., 1997). Collectively, these data have led to the “Pre-Myofibril Model” of sarcomere assembly (Sanger et al., 2005; Sanger et al., 2017). Due to the above mentioned caveats of model systems, whether pre-myofibrils transition into sarcomeres, and the mechanisms governing their formation and transition have remained elusive.

To aid these cell culture studies, a number of groups have also developed *in vivo* systems to study sarcomere assembly. Studies have utilized tissue samples from chick embryos, mice, zebrafish, and genetic *Drosophila* models to investigate sarcomere assembly (Du et al., 2008; Rui et al., 2010; Sanger et al., 2005; Sanger et al., 2017; Sparrow and Schock, 2009). These systems also suffer from important caveats. Namely, a number of model systems, including zebrafish and *Drosophila* investigate skeletal muscle, and not cardiac muscle, which contain vast differences in their function, architecture, and molecular composition(Sanger et al., 2005; Sanger et al., 2017). *In vivo* chick studies suffer from the aforementioned lack of genetic tools. Finally, most *in vivo* work from chick and mice have suffered from a lack of both high temporal and spatial resolution, compared to temporal and spatial resolution capable in cell culture systems. Collectively, *in vivo* model systems has resulted in the lack of a reproducible, testable, quantifiable, and easy to use system with which to investigate cardiac sarcomere assembly.

Despite these drawbacks, *in vivo* investigations have largely yielded descriptive and static evidence to support the Pre-Myofibril Model, but a number of studies have added mechanistic insight and even led to new models of sarcomere assembly (Ma et al., 2009; Rui et al., 2010; Sanger et al., 2005; Sanger et al., 2017; Tullio et al., 1997). For example, beautiful microscopic investigations in chick heart tissue has essentially recapitulated every aspect of cell culture studies, showing the same structures described in the pre-myofibril model exist *in vivo*, specifically localizing NMIIB containing stress fibers adjacent to myofibrils (Du et al., 2008). However, other studies have failed to localize NMIIA or NMIIB containing stress fibers within cardiomyocytes in mouse or chick tissue (Ehler et al., 1999; Kan et al., 2012; Ma et al., 2009). Furthermore, studies using mouse genetics have attempted to investigate the role of both NMIIA and NMIIB. Germline NMIIA knockout (KO) mice fail to gastrulate (an event which precedes heart formation), and a conditional p9 KO NMIIA mouse appears to have no effect on heart formation (Conti et al., 2004; Conti et al., 2015). NMIIB germline KO mice were reported to have disorganized but assembled sarcomeres via EM, though this was never quantified (Tullio et al., 1997). In addition, a conditional heart KO of NMIIB also displays striated sarcomere structures (Ma et al., 2009). Collectively, these studies have called into question the role of NMIIA and NMIIB during sarcomere assembly, and the relevance of pre-myofibril structures.

Work in mice has also implicated the actin filament nucleating formin FHOD3 as important for sarcomere assembly and/or maintenance. Germline KO FHOD3 mice display reduced sarcomeres at p13.5 compared to control animals, suggesting they have sarcomere formation defects (Kan et al., 2012). However, these animals also display a heartbeat at p7 which suggests these animals may have assembled sarcomeres, and instead these animals suffer from a defect in sarcomere maintenance/homeostasis (Kan et al., 2012). Indeed, cell culture studies using neo-natal rat ventricular myocytes have shown that cardiomyocytes lose their sarcomeres after FHOD3 is knocked down (Iskratsch et al., 2010; Taniguchi et al., 2009). Thus, FHOD3 plays a clear role in sarcomere maintenance but its potential role in sarcomere assembly is unclear.

Studies using the body wall (i.e., skeletal muscle) and primary muscle cells of *Drosophila,* chick cardiomyocytes, and BC3H muscle cells have described the presence of “I-Z-I” bodies, composed of thin actin filaments and α-actinin Z-bodies, and “floating” A-bands not containing detectable actin (Holtzer et al., 1997; Lu et al., 1992; Rui et al., 2010). These structures are proposed to be stitched or brought together to form mature sarcomeres and myofibrils eventually containing other sarcomeric proteins (Rui et al., 2010; Sanger et al., 2005). In support of this model, studies in *Drosophila* have shown the presence of small myosin filaments following knockdown (KD) of separate “I-Z-I” components (Rui et al., 2010). Despite the descriptive work showing the presence of I-Z-I bodies in skeletal muscle sources, an I-Z-I complex in cardiac tissue as depicted in models has never been localized in cardiomyocytes, and further, actual stitching or bringing together of I-Z-I or I-Z-I-like bodies and “floating” A-bands has never been shown. While this is conceptually an intriguing model, stitching of either I-Z-I bodies or myosin II filaments has not been observed.

The above discrepancies represent a major roadblock in our understanding of sarcomere assembly. Thus, we sought to test how sarcomeres are assembled using live cells and recent technological advances. Here, we leverage our discovery that immature <u>h</u>uman <u>i</u>nduced pluripotent stem cell derived <u>c</u>ardio<u>m</u>yocytes (hiCMs) completely disassemble and then reassemble their sarcomeres following plating, which serves as an assay of synchronized sarcomere assembly. Using this assay we show that sarcomeres arise *directly* from muscle stress fibers (MSFs). We then define the mechanisms governing the MSF to sarcomere transition. Intriguingly, we show the formation of MSFs and their subsequent transition into sarcomeres are based on mechanisms distinct from those driving the assembly of non-muscle stress fibers. Importantly, our data does not support any one previously proposed model of sarcomere assembly, but, rather some aspects of several models. As such, we now propose a unified model of sarcomere assembly based on mechanisms underlying the formation of MSF and their subsequent transition into sarcomere containing myofibrils.

## Results

### Development of an assay to test *de novo* sarcomere assembly

To address how cardiac sarcomeres are assembled, we used hiCMs as a model system (see **Materials and Methods**) (Takahashi et al., 2007). We first noted the actin in hiCMs, which had spread for 24hrs post plating, had two distinct organizations, muscle stress fibers (MSFs) and sarcomere containing myofibrils (Figure 1B). Spread hiCMs displayed MSFs at the leading edge, and organized sarcomere structures in the cell body (Figure 1B). Strikingly, superresolution imaging revealed the MSFs in hiCMs resembled a classic actin stress fiber found in non-muscle cells, referred to as actin arcs (Figures 1C and 1D) (Heath, 1983; Hotulainen and Lappalainen, 2006). These MSFs in hiCMs also resemble on a physical level, an actin based stress fiber population found in avian cardiomyocytes proposed to template sarcomere containing myofibrils (Rhee et al., 1994). Importantly, the concept that an actin based stress fiber templates the formation of a myofibril has never been experimentally demonstrated.

To test whether MSFs are a template for sarcomere assembly, we first needed to develop an assay of *de novo* sarcomere assembly. Importantly, we noticed that freshly plated hiCMs contained no sarcomere structures (Figure S1). hiCMs subsequently assembled sarcomere structures over the course of 24 hours, with the first sarcomeres appearing at 7.6 +/- 2.1 hours (N=15 hiCMs from 3 independent experiments). Loss of sarcomere structure was confirmed by using multiple sarcomeric, including actin, beta cardiac myosin II (βCMII), *α*-actinin, and TroponinT (Figure S1). We also note that we did not detect the presence of any I-Z-I bodies as presented in representations of the “Stitching Model” (Sanger et al., 2005). As this assay represents a powerful new experimental paradigm to study *de novo* sarcomere assembly, we next sought to directly demonstrate that MSFs template sarcomeres. Indeed, time-lapse microscopy of hiCMs expressing the actin probe Lifeact-mEmerald (Riedl et al., 2008) clearly revealed that MSFs acquire sarcomeres (Figures 1E and 1F). We next wanted to define the mechanisms governing MSFs and their acquisition of sarcomeres. As the mechanisms of actin arc formation and maintenance have been well studied, we were interested in using our assay to test whether the same mechanisms driving actin arc dynamics were governing MSF dynamics (Burnette et al., 2014; Hotulainen and Lappalainen, 2006; Murugesan et al., 2016).

### Actin Retrograde flow in hiCMs and non-muscle cells

Actin arc stress fibers in non-muscle cells undergo robust “retrograde flow” away from the edge of the cell (Forscher and Smith, 1988; Hotulainen and Lappalainen, 2006; Wang, 1985). We confirmed this in a classic model of mesenchymal migration, U2-OS cells (Figures 1G and 1I) (Hotulainen and Lappalainen, 2006). Kymography measurements found that actin arcs in U2-OS cells moved at ~200 nm/min, in agreement with previously published findings (Figures 1G and 1I) (Ponti et al., 2004). We found MSFs also underwent retrograde flow (Figures 1H and 1I). Strikingly however, kymography revealed MSFs moved significantly slower than actin arcs in U2-OS cells (Figures 1G, 1H, and 1I). This was the first indication that actin arcs in non-muscle cells are different that MSFs in hiCMs.

### Formins but not the Arp2/3 complex are required for MSF-based sarcomere formation

The Arp2/3 complex is well known to be required for actin arc formation in non-muscle cells (Hotulainen and Lappalainen, 2006). To test the role of the Arp2/3 complex during sarcomere assembly, we allowed hiCMs to spread in the presence of CK666, an inhibitor of the Arp2/3 complex (Nolen et al., 2009). Surprisingly, hiCMs allowed to spread in the presence of CK666 formed robust MSFs and sarcomeres comparable to untreated control cells (Figures 2A, inset, and 2B). To confirm inhibition of the Arp2/3 complex by CK666, we examined the localization of the Arp2/3 complex with and without CK666 treatment. Importantly, the strong localization of the Arp2/3 complex at the edge of control hiCMs was absent in CK666 treated hiCMs (Figures 2C and 2D). The delocalization of the Arp2/3 complex from the leading edge is consistent with inactivation by CK666, as shown previously in non-muscle cells (Henson et al., 2015).

**Figure 2:**
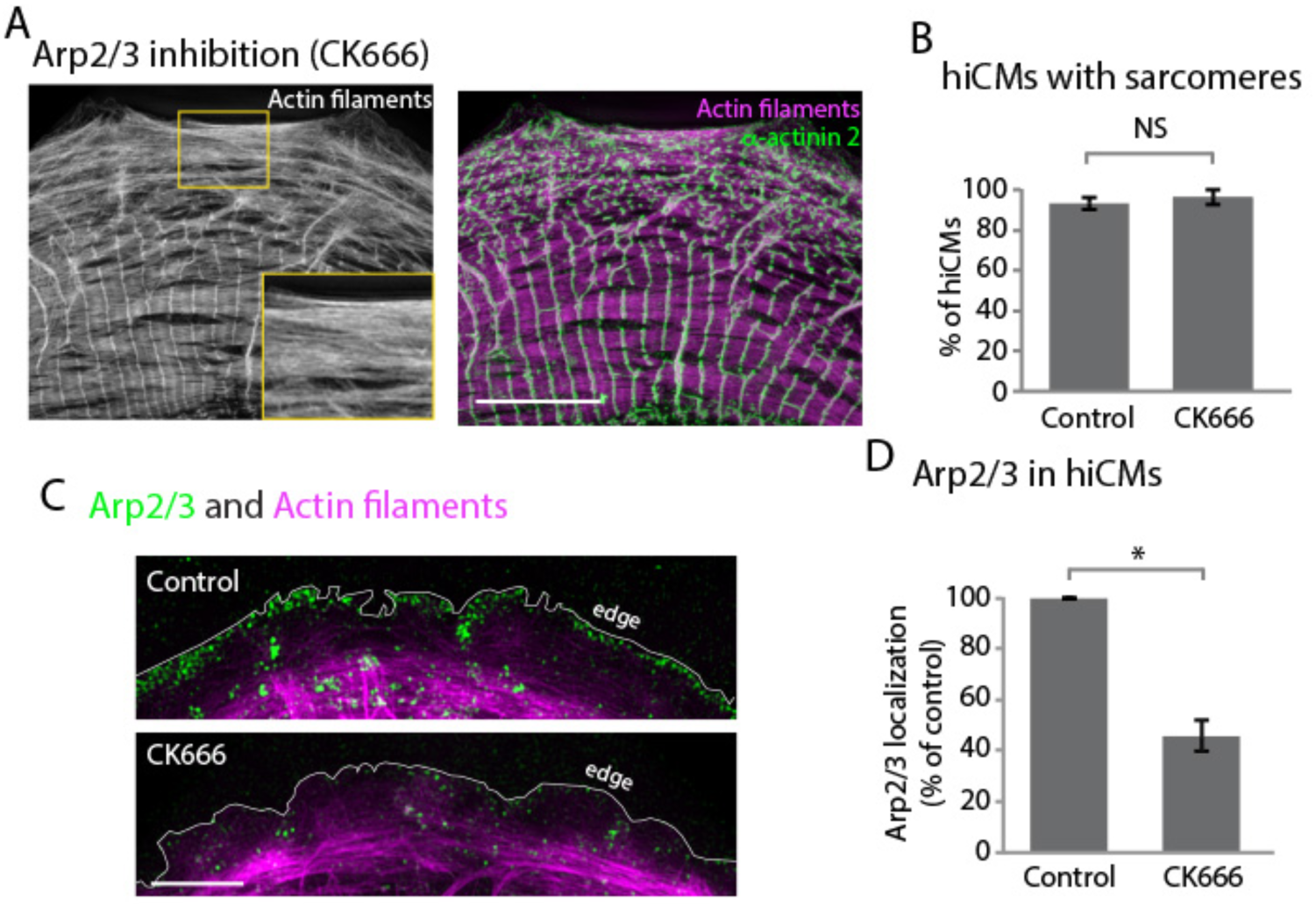
The Arp2/3 complex is not required for sarcomere assembly. (A) hiCM allowed to spread in the presence of 25 μM CK666 and labeled with actin and α-actinin 2 (i.e., Z-lines). Box indicates presence of MSFs. (B) Quantification of percentage of cells with sarcomeres at 24hrs post plating in control and 25 μM CK666. Control: 76 cells, 10 experiments; 25 μM CK666: 41 cells, 3 experiments. (C) 25 μM CK666 treatment results in delocalization of the Arp2/3 complex from the leading edge of hiCM (bottom) compared to untreated controls (top). (D) Quantification of loss of the Arp2/3 complex form the leading edge of hiCMs. Control; 36 cells over 3 experiments. 25 uM CK666; 29 cells over 3 experiments. Scale bars; (A), (C) 10 μm.

In addition to the Arp2/3 complex, formin-mediated actin polymerization has been shown to be crucial for actin arc formation and dynamics in a multitude of cell lines (Hotulainen and Lappalainen, 2006; Murugesan et al., 2016). As a starting point to test whether formins are required for sarcomere assembly, we allowed hiCMs to spread in the presence of a pan-inhibitor of formin-mediated actin polymerization, <u>s</u>mall <u>m</u>olecule Inhibitor of formin homology domain <u>2</u>, SMIFH2 (Rizvi et al., 2009). SMIFH2 has been shown to stop formin mediated actin polymerization, actin arc formation and retrograde flow in cells (Henson et al., 2015; Murugesan et al., 2016; Rizvi et al., 2009). We found hiCMs spreading in the presence of SMIFH2 completely failed to form sarcomeres (Figures 3A and 3B). These results suggest formins are required for *de novo* sarcomere assembly. As there are 15 mammalian formin genes, we next asked what specific formin was required for sarcomere assembly.

Previous data from isolated rat cardiomyocytes has definitively shown FHOD3 as crucial for sarcomere maintenance (Iskratsch et al., 2010; Kan et al., 2012; Taniguchi et al., 2009). These results clearly showed rat cardiomyocytes containing myofibrils subsequently lost their myofibrils following FHOD3 knockdown (Iskratsch et al., 2010; Taniguchi et al., 2009). Despite this definitive result, the role of FHOD3 during sarcomere assembly has not been tested. Therefore, we sought to use our assay to directly test if the formin FHOD3 was required for MSF based sarcomere assembly. Thus, we treated hiCMs with siRNA against FHOD3. Following knockdown (KD) of FHOD3, hiCMs were unable to assemble sarcomeres following plating (Figures 3C and 3D). Indeed, the actin organization in FHOD3 KD hiCMs highly resembled hiCMs spread in the presence of SMIFH2 (Figures 3A and 3C). This data demonstrates FHOD3 is required for *de novo* sarcomere assembly. While this data is consistent with a role of FHOD3 in sarcomere assembly, it does not reveal insight into whether the formin is acting through the MSF, sarcomeres, or both.

We next asked if formin inhibition was affecting either the MSF or sarcomeres directly. To test this, we allowed hiCMs to spread for 24hrs (after they have established sarcomeres) and imaged their actin cytoskeleton via live-cell microscopy before and after administering SMIFH2 (Figure 3E). Following addition of SMIFH2, formation of new MSFs was immediately blocked, along with retrograde flow of existing MSFs (Figures 3F and 3G). However, we did not detect any changes in sarcomere structure, and hiCMs continued to beat in the presence of SMIFH2 (Note sarcomere structure in Figure 3E). Taken together, our data shows sarcomeres derive from MSFs and suggest a crucial role for formin-mediated actin polymerization during sarcomere formation in hiCMs, but not the Arp2/3 complex.

**Figure 3:**
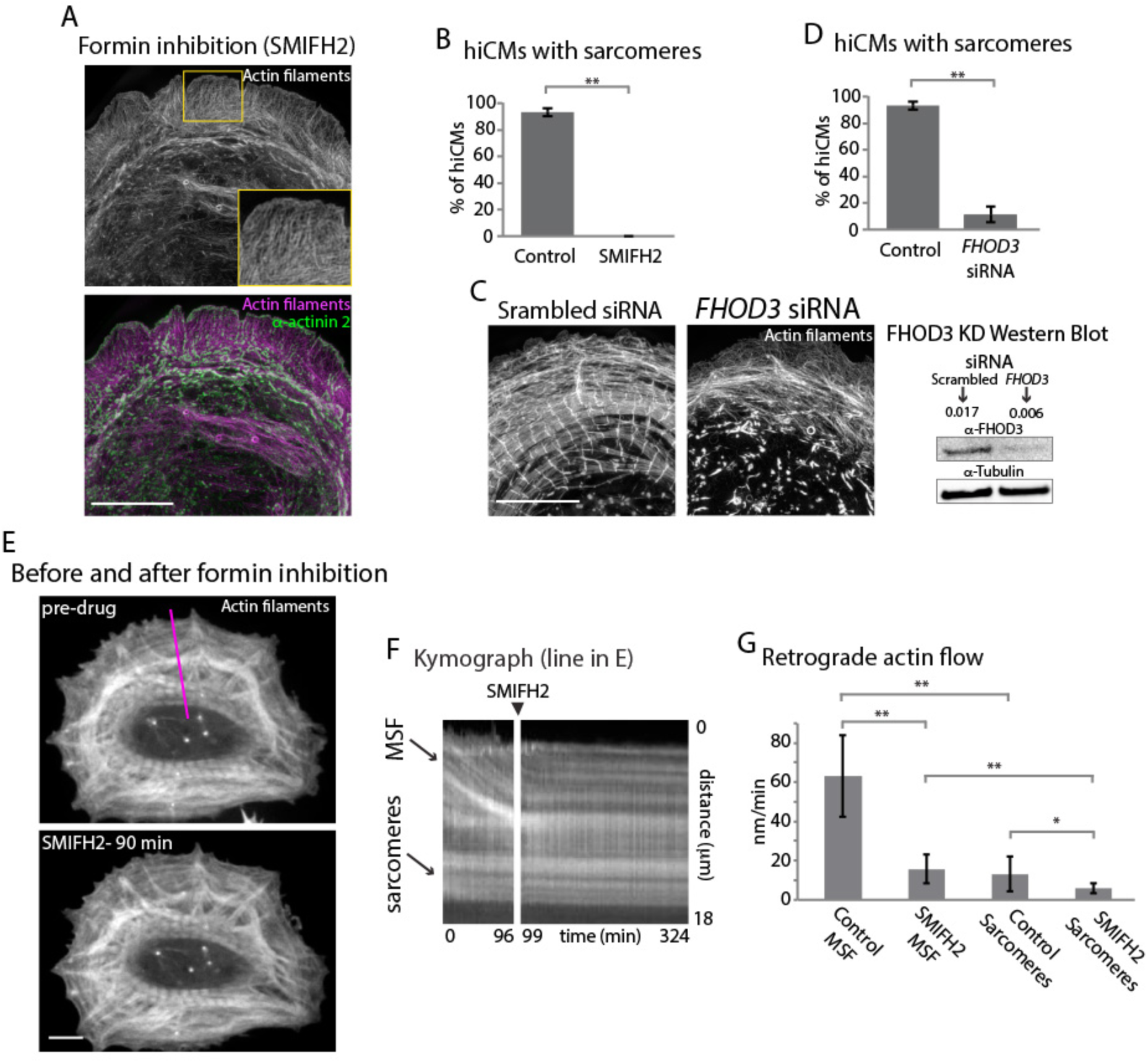
FHOD3 is required for sarcomere assembly and MSF dynamics. (A) hiCM allowed to spread in the presence of 25 μM SMIFH2 and labeled with actin and α-actinin 2. Box indicates loss of transverse MSFs behind leading edge of hiCM. (B) Quantification of percentage of cells with sarcomeres at 24 hrs post plating. (C) siRNA scramble control and siRNA FHOD3 hiCMs allowed to spread for 24 hrs. Note loss of sarcomeres comparable to SMIFH2 treatment in FHOD3 KD hiCMs (Figures 3A and 3B). Western blot (right) denotes protein loss in FHOD3 knockdown. (D) Quantification of percentage of cells with sarcomeres at 24 hrs post plating in control and FHOD3 siRNA conditions. For (B) and (D) quantifications, Control: 76 cells, 10 experiments; 25 μM SMIFH2, 16 cells, 3 experiments; siRNA FHOD3: 33 cells, 3 experiments. (E) Stills from live hiCM expressing Lifeact-mApple before (above) and 90 mins following addition of 25 μM SMIFH2 (below). Note how sarcomeres and overall actin architecture remains unperturbed at 90 mins post 25 μM SMIFH2. (F) Kymograph from purple line in Figure 3E displaying distance over time of MSFs and sarcomeres in hiCM. Left kymograph represents pre-drug control, and right kymograph after addition of 25 μM SMIFH2. Note almost immediate block of retrograde flow of MSFs. (G) MSF and sarcomere retrograde flow rates in hiCMs. Measurements were taken from the leading edge of the cell (for MSFs) and the cell body (for sarcomeres). Control: 12 cells, 3 measurements from each cell, 3 experiments; 25 μM SMIFH2, 12 cells, 3 measurements from each cell, 3 experiments. Scale bars; (A), (C) 10 μm, (E) 5 μm

### NMIIB is required for cardiac sarcomere actin filaments *in vitro* and *in vivo*

In addition to actin nucleators, <u>n</u>on-muscle <u>m</u>yosin <u>II</u> <u>A</u> (NMIIA) activity has been shown to be required for actin arc formation and organization in diverse non-muscle cell types (Burnette et al., 2014; Fenix et al., 2016). Thus, we asked if NMIIA was required for MSF formation and/or the MSF to sarcomere transition. We first localized the two major paralogs of NMII in humans, NMIIA and NMIIB, in spread hiCMs (Vicente-Manzanares et al., 2009). As opposed to nonmuscle cells, which display a distinct distribution of NMII paralogs, with NMIIB being restricted from the leading edge of cells (Kolega, 1998), we found both NMIIA and NMIIB localized to the edge of spreading hiCMs with MSFs, and were restricted from the middle of the cell where the sarcomeres were localized (Figures 4A and 4B). We confirmed the relative pattern of NMII filaments and sarcomeres by immune-localizing NMIIB from P3 mouse heart ventricle tissue. Indeed, we found NMIIB filaments were adjacent to sarcomeres in the mouse heart as it is in hiCMs (Figure S2).

The similar localization patterns of NMIIA and NMIIB led us to ask whether these NMII filaments were in NMII “co-filaments”, a recently reported NMII species that includes both NMIIA and NMIIB paralogs in the same filament (Beach et al., 2014; Shutova et al., 2014). Indeed, super-resolution microscopy revealed that more than 90% of NMII filaments were contained both NMIIA and NMIIB (Figures 4C and S3). These NMII co-filaments highly resemble the cofilaments recently described in non-muscle cells (Beach et al., 2014; Shutova et al., 2014). Indeed, when we measured the lengths of NMII co-filaments in hiCMs, their lengths agreed with previously published measurements (Figure 4D) (Beach et al., 2014; Shutova et al., 2014).

**Figure 4:**
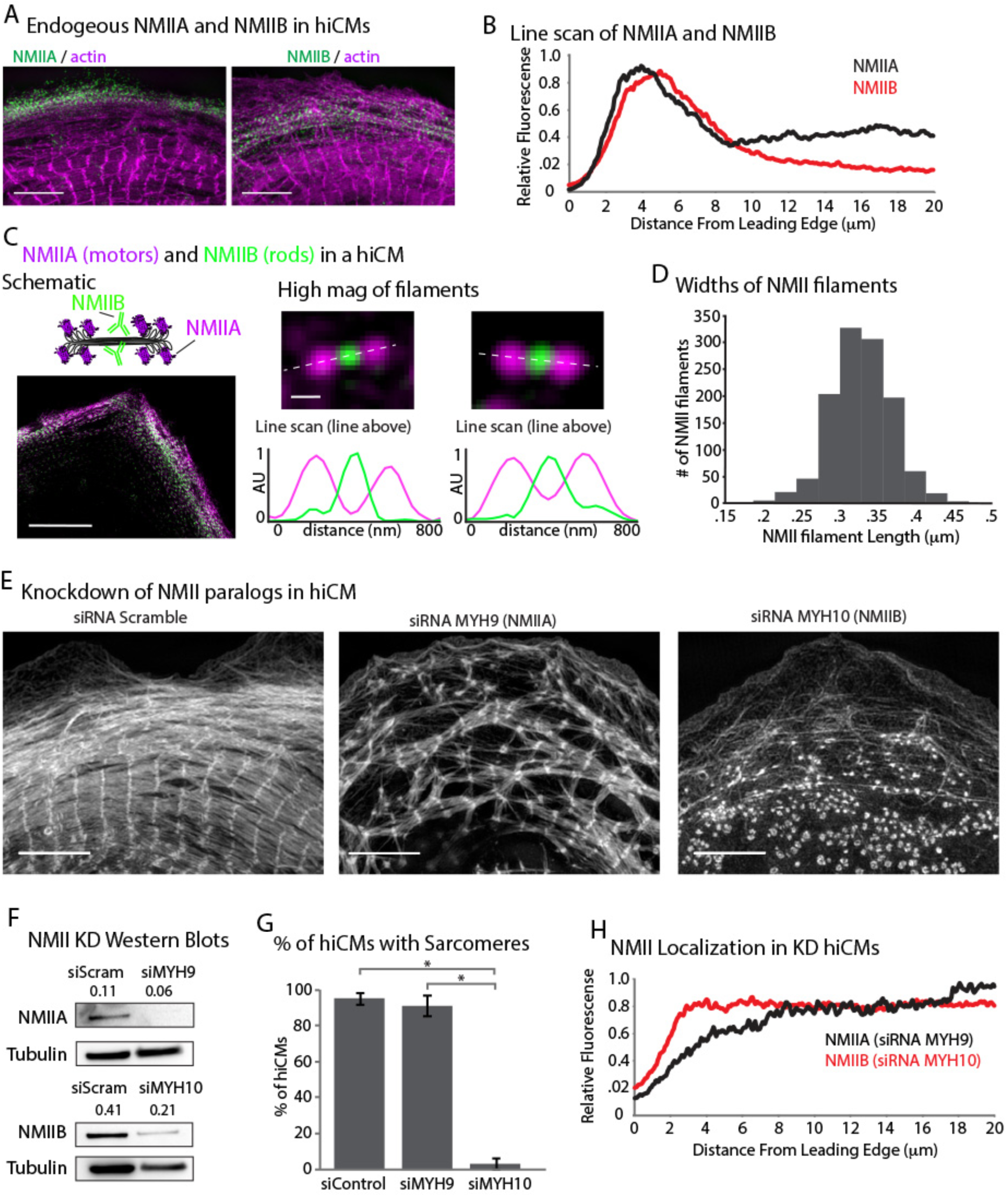
Non-muscle myosin IIB (NMIIB) is required for sarcomere formation. (A) SIM showing NMIIA (left) and NMIIB (right) localized to the edge of hiCMs with MSFs. (B) Line-scans starting from edge of hiCMs showing peak localizations of NMIIA (black) and NMIIB (red) NMIIA: 15 cells, 2 experiments; NMIIB: 32 cells, 4 experiments. (C) hiCM transfected with NMIIA-mEmerald (N-terminal motors) and stained for endogenous NMIIB (left). High-mag views of NMIIA-NMIIB co-filaments (right, top). Line scans across NMII co-filaments above, from N-terminal motors (purple) and C-terminal rod domains (green). (D) Quantification of NMII cofilament length. Histogram displays the distribution of NMII co-filament lengths (motor-domain to motor-domain) NUMBER OF FILAMENTS. (E) Actin of representative control, NMIIA KD (siRNA MYH9), and NMIIB KD (siRNA MYH10) hiCMs allowed to spread for 24hrs. NMIIA KD hiCMs display disorganized sarcomeres, while NMIIB KD hiCMs display no actin-based sarcomeres. (F) Representative western blots of 2 experiments showing knockdown of NMIIA (siRNA MYH9) and NMIIB (siRNA MYH10). (G) Percentage of control, NMIIA KD (siRNA MYH9), and NMIIB KD (siRNA MYH10) hiCMs with actin based sarcomeres at 24hrs spread. Control: 49 cells, 6 experiments; NMIIA KD: 34 cells, 3 experiments; NMIIB KD: 59 cells, 4 experiments. (H) Line-scans of NMIIA and NMIIB from NMIIA (MYH9 KD) and NMIIB (MYH10 KD) KD hiCMs respectively. NMIIA KD: 16 cells 2 experiments; NMIIB KD: 42 cells, 4 experiments. Note how both NMIIA and NMIIB KD cells lose peak localization at leading edge. Scale bars; (A) 5 μm, (C) left 10 μm, (C) right 200 nm, (E) 5 μm.

Given the presence of both NMIIA and NMIIB in each filament within MSFs, we next asked whether NMIIA and/or NMIIB were required for sarcomere assembly. We have previously shown that KD of NMIIA results in loss of actin arcs in U2-OS cells as well as absolute loss of NMIIB filaments (Burnette et al., 2014; Fenix et al., 2016). Thus we hypothesized that NMIIA would likely be the key paralog required for MSF formation and subsequent sarcomere assembly. Surprisingly, KD of NMIIA did not result in an inhibition of MSF or *de novo* sarcomere assembly (Figures 4E to 4G), although the sarcomeres in NMIIA KD hiCMs were disorganized (Figures 4E). Taken together, this data suggests that NMIIA plays an organizational role but is not formally required for sarcomere assembly in hiCMs.

We next depleted hiCMs of NMIIB to test if it was required for MSF formation and sarcomere assembly. Although NMIIB is not required for actin arc assembly in non-muscle cells (Figure S4), its KD resulted in a complete inability of hiCMs to form sarcomeres *de novo* after plating (Figures 4E to 4G). These results argue NMIIB is the major paralog required for sarcomere assembly in hiCMs. This is opposed to non-muscle cells, where NMIIA has been implicated as the NMII paralog responsible for actin arc stress fiber formation and dynamics (Fenix et al., 2016). Live-cell microscopy revealed NMIIB KD hiCMs displayed dramatic defects in actin retrograde flow, and increased protrusion/retraction events compared to control siRNA treated hiCMs (Supplemental Movies 1 and 2). While these defects in sarcomere assembly were dramatic, we noticed that hiCMs treated with siRNA against NMIIB were still beating before replating (Supplemental Movies 3 and 4). This implied that the pre-existing sarcomeres of the hiCMs were still intact after KD. Therefore, we immuno-localized α-actinin 2 to visualize sarcomeres. Surprisingly, we found there were no differences in control and NMIIB KD cells before plating (Figures 5A and 5B). Collectively, this data would suggest NMIIB is required for *de novo* sarcomere formation, but not homeostasis (i.e., turnover) of pre-existing sarcomeres.

We next sought to test whether NMIIB was required for sarcomere assembly *in vivo*. Data from previous studies attempting to answer this question in mice have been difficult to interpret. Germline NMIIB KO mice form a functional heart (Tullio et al., 1997). Only one high magnification image of the KO animal was shown, which showed severe sarcomere disorganization (Tullio et al., 1997). Of interest, the NMIIB KO animals also showed highly increased NMIIA protein levels compared to controls (Tullio et al., 1997). Thus, the observed capacity of this animal to form sarcomere like structures could be due to genetic compensation by NMIIA. A conditional KO NMIIB mouse has also been made (Ma et al., 2009). This mouse model has several issues. First, while the authors showed an impressive NMIIB KD in the cerebellum via a neuronal specific driver, there were high levels of NMIIB protein in the heart at the time of analysis in the heart specific KO (Ma et al., 2009). Second, the heart specific conditional KO is driven off of the alpha myosin heavy chain promoter, which switches on *after* sarcomere assembly has begun and indeed after the heart fields have begun to beat (Ma et al., 2009; Ng et al., 1991). While the only high-resolution images focused on the intercalated discs, the presence of sarcomeres is easily observed, though their organization is difficult to assess (Ma et al., 2009). While this mouse model supports our conclusion that NMIIB is not required for the the maintenance of pre-existing sarcomeres, it also demonstrates that the mouse may not be the optimum model to test the role of NMIIB in sarcomere assembly.

We next sought an *in vivo* model where rapid KD of NMIIB could be achieved before sarcomere assembly has begun. In zebrafish, rapid KD of NMIIB can be achieved via morpholino (MO) before heart formation with little genetic compensation as seen in KO animals (Gutzman et al., 2015; Kimmel et al., 1995). Therefore, single cell zebrafish embryos were injected with control MO or *myh10* MO directed against NMIIB as previously described (Gutzman et al., 2015). Embryos were fixed 48 hours post fertilization, actin filaments were labeled, and the lengths of of sarcomeres were quantified. Animals injected with MO directed against NMIIB formed a proper 2 chambered heart. However, *myh10* MO animals had a significant decrease in sarcomeres (Figures 5C and 5D). Taken together, our data shows that KD of NMIIB results in a reduction of sarcomere assembly *in vivo*, which is consistent with our findings in hiCM.

### NMIIB is required for organized A-band formation

Thus far, our results highlight the importance of formin-mediated actin polymerization and NMIIB for proper actin filament architecture during sarcomere assembly. We next wanted to address how the thick, <u>β</u> <u>C</u>ardiac <u>M</u>yosin <u>II</u> (βCMII), filaments at the core of the sarcomere (i.e., A-band, Figure 1A) assemble. Therefore, we started by localizing endogenous βCMII filaments and NMIIB filaments. βCMII predominately localized behind NMIIB in organized sarcomere structures, and showed a peak localization ~15 microns behind the leading edge of the cell, with a slight area of overlap with NMIIB (Figures 6A and 6B). We noticed that near the leading edge of the cell, βCMII filaments were smaller and and not organized into stacks resembling A-bands (Figure 6 C). In addition, we also noted these short βCMII filaments also contained NMIIB filaments, suggesting the presence of βCMII-NMIIB co-filaments (Figures 6C, 6D and S3). To our knowledge, this is the first time a myosin II species has been reported that contains a non-muscle and muscle paralog inside of cells. Furthermore, we also found NMIIB-βCMII cofilaments in mouse and human heart tissue, indicating NMIIB-βCMII co-filaments are present *in vivo* (Figures 6C and S3). It is also important to note that we only detect NMIIB-βCMII co-filaments in hiCM in the region of overlap between the NMIIB and βCMII localizations, where the lengths of βCMII filaments are short and not in longer βCMII filaments further away from the edge (Figure 6D and 6E). The presence in of NMIIB before βCMII filaments grow into larger filaments led us to test the hypothesis that NMIIB would play a role in βCMII filament formation.

**Figure 6:**
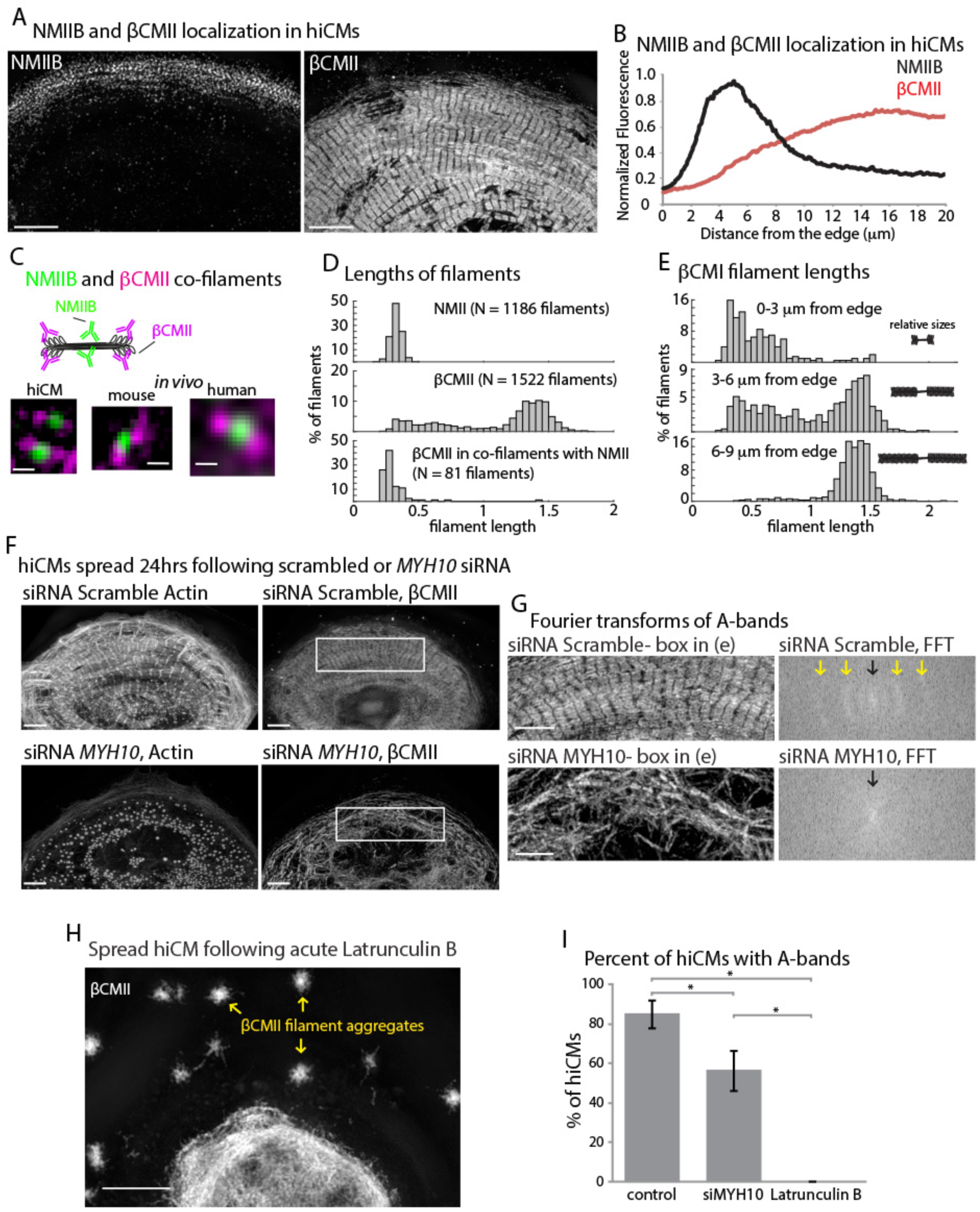
Organized βCMII A-bands require proper actin organization. (A) Endogenous localization of NMIIB (left) and βCMII in hiCMs. (B) Averaged linescans of NMIIB (Figure 4B) and βCMII localization in hiCMs spread for 24hrs. Note the peak fluorescence of βCMII is more towards cell body than peak fluorescence of NMIIB. 23 cells from 4 experiments were used for βCMII localization. (C) Schematic (top) of NMIIB-βCMII filaments in hiCMs. High-mag views of βCMII-NMIIB co-filaments. Endogenous staining of hiCM for βCMII (N-terminal motors) and NMIIB (rod domain). Mouse and human tissue (middle, right) stained for βCMII (motors) and NMIIB (rod domain). (D) Histograms displaying width of NMII filaments (top), βCMII filaments (middle), and NMIIB-βcMlI co-filaments in hiCMs. (E) Histograms displaying distribution of βCMII filaments widths with respect to their location of the cell. Note βCMII filaments tend to grow larger as they move towards the center of the cell. Measurements were not taken from “mature” sarcomere structures in highly organized A-bands. (F) Actin and βCMII of control hiCM spread for 24hrs (above). NMIIB KD (siRNA myh10) hiCM spread for 24hrs (below). Note loss of organized A-bands but presence of βCMII filaments. (G) Fourier transforms of βCMII signal from white boxes in Figure 6F from control and NMIIB KD hiCMs (above and below respectively). Yellow arrows indicate sarcomeric periodicity in control cells, which is lacking in NMIIB KD cells. (H) βCMII in hiCM spread for total of 24hrs, with final 6 hours in 5 μM Latrunculin B. Notice lack of βCMII A-bands, and large βCMII filament aggregates (yellow arrows). (I) Percentage of control, NMIIB KD (siRNA *myh10*), and Latrunculin B hiCMs with βCMII A-bands. Control: 26 cells, 3 experiments; NMIIB KD: 26 cells, 2 experiments; Latrunculin B: 11 cells, 3 experiments. Scale Bars; (A) 5 μm, (C) 200 nm, (F), (G), (H) 5 μm.

To test if NMIIB was also required for βCMII filament and A-band formation we depleted hiCMs of NMIIB and localized βCMII 24hrs after plating. Compared to control hiCMs, NMIIB KD hiCMs displayed a significant decrease in the ability to form A-band like structures, but, surprisingly, retained the ability to form βCMII filaments (Figures 6F to 6I). Although (3CMII filaments formed, they were highly disorganized compared to control cells, as assessed by Fourier transform (Pasqualini et al., 2015) (Figure 6G). As NMIIB KD results in highly disorganized actin filament architecture, we asked if βCMII was using residual actin filaments as a template to polymerize.

To test if actin was serving as a template for βCMII filament formation, we sought to remove all the actin filaments in hiCMs. To do this we allowed hiCMs to form sarcomeres for 18 hrs, and treated hiCMs with the actin monomer sequestration agent, Latrunculin B, for 6 hours in order to disassemble actin filaments acutely before fixation (Spector et al., 1983; Wakatsuki et al., 2001). Latrunculin B treated hiCMs showed a lack of actin based sarcomeres, and no βCMII filament stacks, but conspicuous βCMII filament aggregates (Figures 6H and 6I). These results argue that MSFs and sarcomeric actin filaments are serving as “tracks” with which βCMII filaments are loading onto as they form larger βCMII filament stacks of the A-band.

Although the NMIIB-βCMII co-filaments are of interest, due to their relatively low numbers, we do not believe they are the major mechanism through which NMIIB is facilitating βCMII filament stack assembly (i.e., the A-band). Therefore, we were interested in the physical mechanisms of A-band formation. To address this question, we created a full length, human βCMII construct containing a mEGFP tag on the motor domain (i.e., N-terminal) (Figure 7A). This construct properly integrated into both single filaments and more mature myofibrils (Figures 7A and 7B). We previously showed a similar myosin, NMIIA, primarily formed A-band-like filament stacks through expansion of a single filament or tight bundle of filaments, and to a much lesser degree, concatenation of separate filaments to form a larger ensemble (Fenix et al., 2016). To test how βCMII filament stacks form, we repeated our live-cell sarcomere formation assay using our βCMII-mEGFP construct. In contrast to our previous results with NMIIA, we found the major physical mechanism of βCMII filament stack formation to be concatenation, where pre-existing βCMII filaments ran into and stitched together to form the A-band (Figures 7C and 7E). A small percentage of hiCMs showed an expansion event of βCMII-mEGFP, however this was significantly less frequent than in non-muscle cells, and did not appear to result in a more organized A-band (Figures 7D and 7E). Indeed, each of the hiCMs quantified in Figure 7E showed only 1 expansion event.

**Figure 7:**
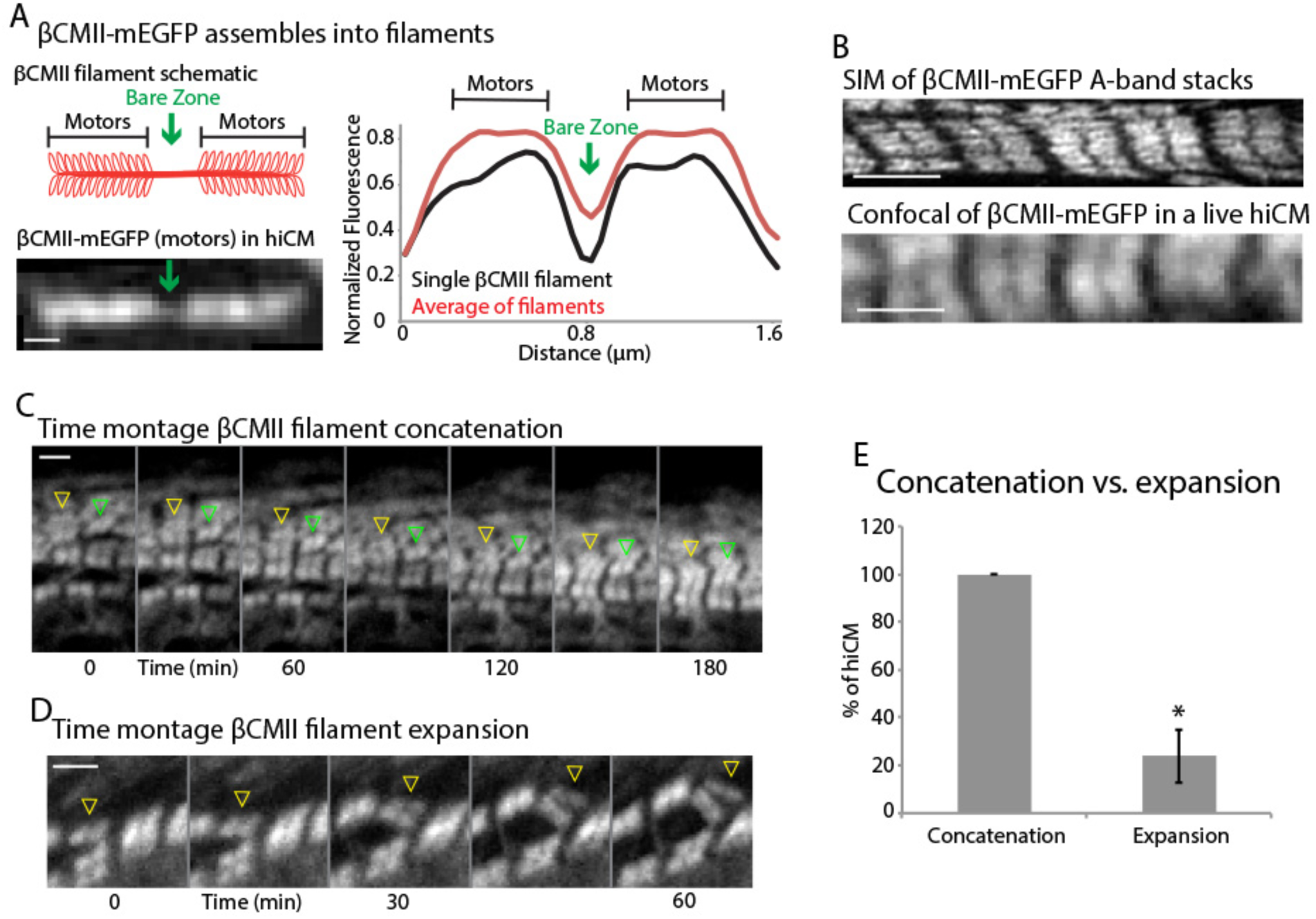
βCMII filaments concatenate to form larger A-band structures. (A) Cartoon of βCMII filament (left, above). N-terminal tagged human βCMII-mEGFP filament expressed in hiCM (below). Gap in signal represents bare-zone lacking motors. (βCMII single filament and A-band filaments (βCMII filaments found within organized A-bands as in Figure 7B) widths measured by line scans (right). βCMII Filaments: 16 filaments, 3 experiments. βCMII myofibrils: 28 myofibrils, 3 experiments. Note more level “plateau” of signal from motors in A-band βCMII filaments. (B) Structured Illumination Microscopy (SIM) of representative βCMII-mEGFP myofibril in hiCM (top) and laser scanning confocal (bottom). (C) Montage of βCMII A-band stack forming via concatenation of separate filaments in hiCM transfected with βCMII-mEGFP (yellow arrows). (C) Representative montage showing two separate concatenation events. Yellow arrowhead denotes a large stack of βCMII filaments concatenating with a smaller stack of βCMII filaments as they undergo retrograde flow. Green arrowhead denotes smaller βCMII filament concatenating with larger βCMII stack as they undergo retrograde flow. Both events result in larger and more organized βCMII filament stack (i.e. the A-band). (D) Example of βCMII filament splitting event. Note how small βCMII stack splits to create 2 smaller βCMII filaments and does not result in larger or more organized βCMII filament stacks. (E) Quantification of % of hiCMs which display concatenation and/or expansion events of βCMII filaments. Scale Bars; (A) 200 nm, (B), (C) and (D) 2 μm.

## Discussion

The purpose of this study was to address a decades-old question concerning the mechanisms underlying actin and myosin based sarcomere assembly. A number of labs have proposed sarcomeres arise from an actin-based precursor. Based on previous actin filament staining and NMII localization to these stress fibers, our initial hypothesis was that the mechanisms underlying actin filament polymerization and organization would be related to those governing the assembly of non-muscle stress fibers. In support of this hypothesis, both previous reports in non-human cardiomyocytes and our own work with hiCMs show that the actin organization of these precursors appears identical to the classic non-muscle stress fiber population known as actin arcs (Heath, 1983; Tojkander et al., 2012). Indeed, both sarcomere precursors and actin arcs display retrograde flow by which they move away from the cell’s edge (Figure 1). However, these were the only similarities that our study revealed. Instead, our data argues that these sarcomere precursors represent an actin stress fiber regulated by different mechanisms than those regulating non-muscle stress fibers. We refer to this sarcomere precursor as a Muscle Stress Fiber (MSF). On a dynamic level, MSFs translocate substantially slower than actin arcs. MSFs also acquired sarcomeres, whereas actin arcs, obviously, did not. Even the actin nucleating factors required for MSF and sarcomere assembly were different.

It has been well established that the actin filaments of actin arcs require both nucleation mediated by the Arp2/3 complex and formins. The Arp2/3 complex does localize to the edge of hiCMs in a similar manner as it does to the edge of non-muscle cells. However, we found that Arp2/3’s removal from the edge by a specific inhibitor did not did not prevent the assembly of MSFs or their transition into sarcomeres (Figure 2). Thus, Arp2/3 appears to be required for the assembly of actin arcs and but not MSFs in hiCMs. The specific role, if any, of the Arp2/3 complex in cardiomyocytes will be of interest in the future. In contrast to the Arp2/3 complex, our data suggests that formins are required for MSFs and sarcomere assembly. Pharmacological inhibition of formins resulted in an ablation of MSF and sarcomeres in hiCMs. On the surface, this appears to be a similarity between actin arcs and MSFs. However, we found that the formin paralog, FHOD3, was responsible for MSF and sarcomere assembly, whereas in non-muscle cells mDia1 is required for actin arcs (Murugesan et al., 2016). This adds to the already established role of FHOD3 in sarcomere homeostasis (i.e., turn-over) (Iskratsch et al., 2010; Kan et al., 2012; Taniguchi et al., 2009). Of note, FHOD3 is expressed in the heart, kidney, and brain, and is the most highly expressed formin in the heart, making it unlikely to be required for the assembly of actin arcs in non-muscle cells (Iskratsch et al., 2010; Kanaya et al., 2005; Katoh and Katoh, 2004). Not surprisingly, we found that U2-OS and HeLa cells did not express FHOD3 and treatment with siRNA directed against FHOD3 in U2OS cells had no effect on actin arcs (data not shown). Taken together, our data shows that the mechanisms underlying actin filament nucleation in MSF are distinct from actin arcs, with significant divergence even with respect to the formin family paralog involved in their formation

Our finding that proper MSF and sarcomere assembly is perturbed by NMIIB KD is surprising. The localization of NMIIB shown by Sanger, Sanger, and colleagues have long implicated a role for this protein. However, the requirement for NMIIB during sarcomere assembly has been tested using knockout mouse models with mixed results. A germline knock-out of NMIIB did result in severe sarcomere defects (Tullio et al., 1997). However, there were still some sarcomeres evident in the single electron micrograph presented. In addition, knockout of NMIIB resulted in genetic compensation by up-regulation of NMIIA (Tullio et al., 1997). Therefore, the results are difficult to interpret. In attempt to bypass this genetic compensation, conditional NMIIB knock-out mice were made (Ma et al., 2009). These experiments were complicated by two major experimental factors beyond the control of the researchers. First, this conditional NMIIB knock-out mouse showed inefficient knockdown of NMIIB protein levels, with more than ~50% of NMIIB protein levels remaining in the heart at time of analysis (no quantifications were provided) (Ma et al., 2009). Secondly and most importantly, the promoter used to drive NMIIB KO (i.e., alpha myosin heavy chain) is turned on only *after* sarcomere assembly has begun in mice (Ma et al., 2009). As our experiments show, NMIIB is required for only *de novo* assembly of sarcomeres, but not their homeostasis (Figure 5A). This means that removing NMIIB after sarcomeres are established should have no effect on their structure (Figure 5A). As such, we find the fact that the conditional NMIIB KO heart muscle appeared to have sarcomeres not surprising. Taken together, the work on mice led us to use acute KD in hiCM versus CRISPR-based knock-out of iPSC’s which would then have to be differentiated over several weeks. Indeed, inhibition of NMII has been shown to increase pluripotency markers in iPSC and embryonic stem cells (Walker et al., 2010).

**Figure 5:**
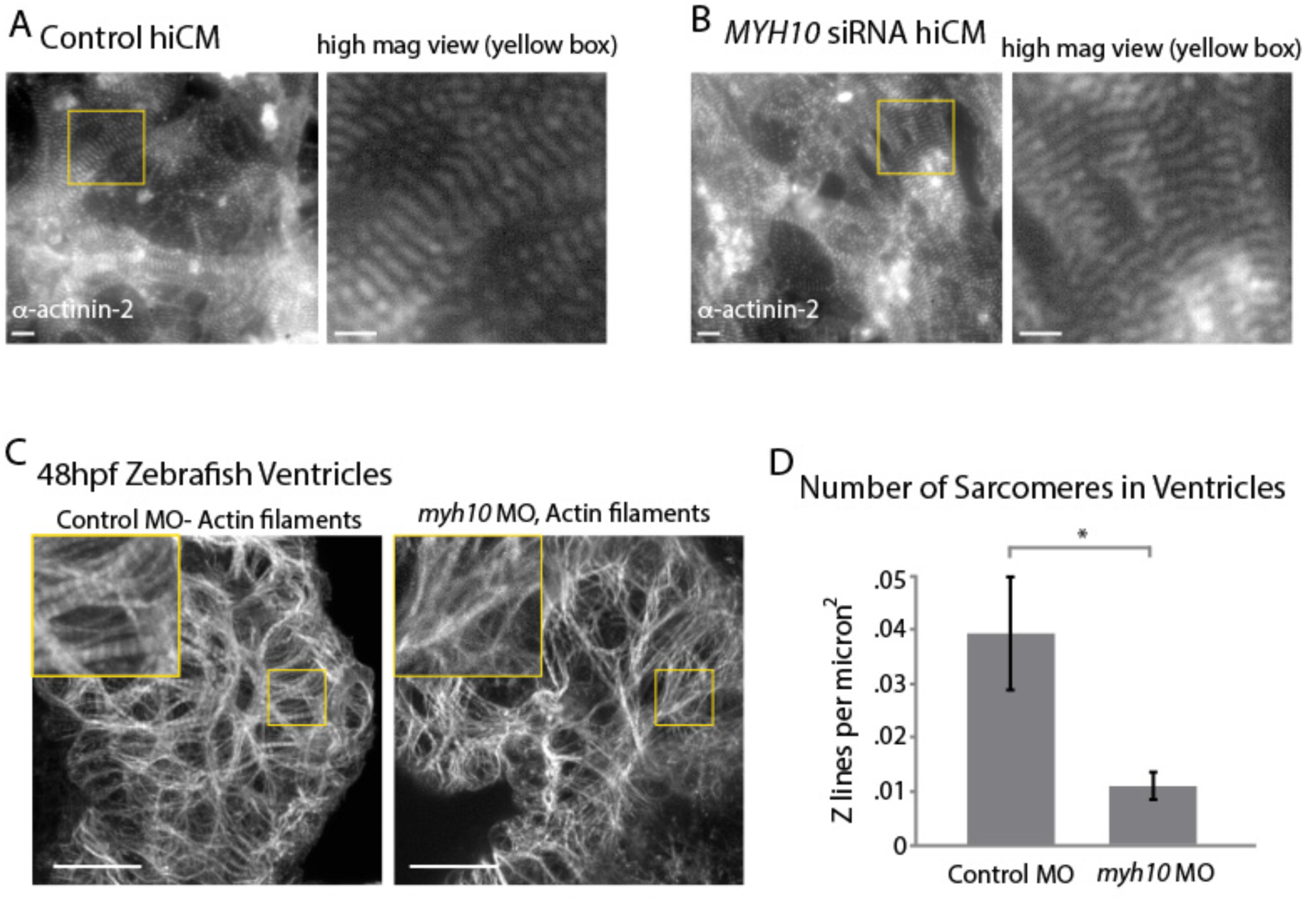
NMIIB is required for *de novo* sarcomere assembly. (A) hiCMs treated with control siRNA in 96 well plate before plating and stained for α-actinin-2 (left). High-mag view (right) showing striated sarcomere structure. (B) hiCMs treated with *MYH10* siRNA in 96 well plate before trypsinization and re-plating and stained for α-actinin-2 (left). High-mag view (right) showing striated sarcomere structure comparable to control cells (Figure 5A). (C) Ventricles of control and *myh10* MO treated zebrafish embryos fixed at 48hrs hpf. Enlarged boxes show myofibrils with sarcomeric pattern in control, which is lacking in *myh10* MO zebrafish. (D) Quantification of number of Z lines/μm in control and *myh10* MO treated zebrafish embryos at 48 hours post fertilization. Control MO: 15 animals, 3 experiments; *myh10* MO: 17 animals, 3 experiments. Scale Bars, A (left), 10 μm; A (right) 5 μm; B (left); 10 μm; B (right), 5 μm; (C) 10 μm.

Genetic compensation after KO of NMII paralogs as well as the need to knock-down NMIIB before sarcomere assembly initiates made us seek a new *in vivo* model system to test the role of NMIIB during cardiac sarcomere assembly. Morpholino-based KD using Zebrafish was the ideal system for this line of experiments. First, it is not a KO that will trigger genetic compensation, the mechanisms of which are not understood at all. Secondly, morpholino induced KD of NMIIB in Zebrafish begins before sarcomere assembly begins, and thus can be used to test *de novo* sarcomere assembly. Indeed, KD of NMIIB in Zebrafish embryos shows a robust decrease in sarcomere assembly compared to control. This data agrees with our *in vitro* data from hiCMs showing NMIIB is required for *de novo* sarcomere assembly.

Thus far, our data underlying how actin filaments of sarcomeres assemble agrees with the Pre-myofibril model proposed by Sanger and Sanger. Indeed, we show sarcomeres arise from a stress fiber precursor (i.e., MSFs) which undergoes a transition as it moves away from the cell edge. However, the Pre-myofibril model has been presented as a mutually-exclusive model to models preceding it, known as the Templating and Stitching models. Our data on how the thick βCMII filaments of the A-band assemble is more in line with the Templating and Stitching models. We found in hiCMs, that βCMII filaments range in size from ~300 nm to ~1.6 microns. βCMII filaments appeared to grow larger the further they moved from the edge of the cell. Interestingly, the smaller βCMII filaments closer to the edge were in myosin II co-filaments with NMIIB (Figures 6C and S3). Given that NMIIB filaments are in co-filaments with βCMII filaments, but not in more mature sarcomeric A-bands, this data would agree with aspects of the Templating model. Specifically, on a molecular level, NMIIB filaments may be serving as a template to seed the formation of βCMII filaments.

Using live-cell microscopy, we show fully formed, separate βCMII filaments concatenate and clearly stitch together to form the A-band. This would agree with aspects of the previously proposed Stitching Model. Specifically, this model proposed individually formed units of the sarcomere (such as myosin II filaments, and so called “I-Z-I bodies”) are brought and subsequently “stitched” together to form myofibrils. While we don’t detect I-Z-I bodies our data clearly shows βCMII filaments stitch together to form the A-band.

Taken together, our data does not support in full any previously proposed model of sarcomere assembly. Instead, our data supports aspects of multiple of these aforementioned models. In our opinion, considering each model of sarcomere assembly as mutually exclusive going forward is not a useful paradigm. Rather, this is conceptually and experimentally limiting. Thus, we present a new and unified model of sarcomere assembly which contains aspects of previously proposed models which we have now experimentally tested (Graphical Abstract and Figure S5). In addition to delineating mechanisms of sarcomere assembly, our model highlights key differences between non-muscle stress fiber and MSF molecular organization, regulation, and dynamics. Simply considering precursor structures as MSFs which transition to a sarcomere containing structure (i.e., myofibril) through a variety of mechanisms will be the most useful model going forward. We refer to this new model as the Muscle Stress Fiber Model of Sarcomere Assembly.

## Methods

### CONTACT FOR REAGENT AND RESOURCE SHARING

*Further information and requires for resources and reagents should be directed to and will be fulfilled by the Lead Contact, Dylan T. Burnette*

### EXPERIMENTAL MODEL AND SUBJECT DETAILS

#### Cell Line Growth and Experimental Conditions

Human induced pluripotent stem cell cardiomyocytes (hiCMs) were purchased from both Axiogenesis (Cor.4u, Cologne, Axiogenesis, now Ncardia) and Cellular Dynamics International (iCell caridomyocytes^2^, Madison, WI). Cells were cultured as per manufacturer’s instructions. hiCMs were cultured in 96 well plates and maintained in proprietary manufacturer provided media until ready for experimental use. Knockdown experiments in hiCMs were started 4 days after the initial thaw. See **Plating Assay** for more detailed protocol for cell plating. See below for detailed information on Cellular Dynamics and Axiogenesis protein transfection and siRNA mediated knockdown

U-2 OS (HTB-96; American Type Culture Collection, Manassas, VA) cells were cultured in 25-cm^2^ cell culture flasks (25-207; Genessee Scientific Corporation, San Diego, CA) with growth medium comprising DMEM (10-013-CV; Mediatech, Manassas, VA) containing 4.5 g/L L-glutamine, D-glucose, and sodium pyruvate and supplemented with 10% fetal bovine serum (F2442; Sigma-Aldrich, St. Louis, MO). For protein expression experiments in U-2 OS cells, cells were transiently transfected with FuGENE 6 (E2691; Promega, Madison, WI) according to the manufacturer’s instructions. Knockdown of NMIIA and NMIIB was performed as previously, with the Accell SMARTpool siRNA to human MYH9, MYH10 or Accell scrambled control purchased from Thermo Fisher Scientific (Waltham, MA) in combination with Lipofectamine 2000 Transfection Reagent (cat# 11668027, Thermo Fisher Scientific, Waltham, MA) for HeLa cells. For both live and fixed cell microscopy, cells were plated and imaged on 35 mm glass bottom dishes with a 10mm micro-well #1.5 cover glass (Cellvis, Mountain View, CA) coated with 25 μg/mL laminin (114956-81-9, Sigma-Aldrich).

#### Animal Conditions

##### Zebrafish maintenance and Husbandry

Standard procedures were used to raise and maintain zebrafish (Kimmel et al., 1995; Westerfield, 2000). All experiments were conducted using the wild-type AB strain zebrafish in accordance with the University of Wisconsin-Milwaukee Institutional Animal Care and Use Committee.

### METHOD DETAILS

#### Plasmids

Plasmids encoding NMIIA-(N-terminal)-mEGFP (11347; Addgene, Cambridge, MA) with mEGFP on the N-terminus of NMIIA heavy chain were used as described previously (Chua et al., 2009). Plasmid encoding Lifeact-mEmerald and Lifeact-mApple were gifts from Michael Davidson. Plasmid encoding NMIIB-(N-terminal)-mEmerald was purchased from Addgene (54192; Addgene, Cambridge, MA). Plasmid encoding human βCMII was synthesized by Genscript (Piscataway, NJ, USA). Briefly, the wild-type human MYH7 (βCMII) sequence from the National Center for Biotechnology Information (NCBI, Bethesda, MD, USA) was cloned into a pUC57 along with Gateway DNA recombination sequences in order to facilitate rapid fluorescent protein integration and swapping (Gateway Technology, ThermoFisher Scientific, Waltham, MA). mEGFP containing a previously published linker sequence was added to the βCMII plasmid using Gateway Vector Conversion System with One Shot *ccdB* Survival Cells (ThermoFisher Scientific, Waltham, Ma) for the βCMII-(N-terminal)-mEGFP construct used in this study.

#### Live-Cell Sarcomere Assembly Visualization Assay

To visualize sarcomere formation, we developed a repeatable method which can be used to visualize any fluorescently tagged protein during sarcomere formation in hiCMs. Prior to performing this assay, cells are maintained in desired culture vessel (our hiCMs were maintained in 96 well plates). In this assay, cells are transiently transfected with desired fluorescently tagged protein, trypsinized, plated on desired imaging dish, and imaged to observe sarcomere formation. A sarcomere assembly assay proceeds as follows. First, hiCMs are transfected with desired fluorescently tagged protein as described below (Viafect, overnight transfection). Transfection mix is washed out with culture media. Cells are then detached from culture vessel using trypsinization method described below. Cells are then plated on a desired culture vessel (in this study, 10mm #1.5 glass bottom dishes, CellVis, were used) for live cell imaging. Imaging vessels were pre-coated with 25 μg/mL laminin for 1 hour at 37° C and washed with 1x PBS containing no Mg^2+^/Ca^2+^. Cells were allowed to attach for ~1.5hrs and media was added to fill the glass bottom dish. Cells were then imaged using either wide-field or laser-scanning microscopy at 3 hrs post plating. The 3 hr time point was optimal, as cells had not yet established sarcomeres, but were healthy enough to tolerate fluorescence imaging. Imaging conditions (e.g., exposure, time intervals, sectioning, etc..) were then modified for each particular probe. This powerful assay can be adapted to visualize the kinetics during sarcomere formation of any fluorescently tagged protein. We have used this assay to visualize actin (Lifeact), βCMII, NMIIA, and NMIIB. The latter three proteins being large, >250 KD proteins.

#### Trypsinization of hiCMs

To trypsinize Cellular Dynamics hiCMs, manufacturers recommendations were used, as follows. All volumes apply were modified from 24 well format for 96 well plates. hiCMs were washed 2x with 100uL 1x PBS with no Ca^2+^/Mg^2+^ (PBS*). PBS* was completely removed from hiCMs and 40uL 0.1% Trypsin-EDTA with no phenol red (Invitrogen) was added to hiCMs and placed at 37° C for 2 minutes. Following incubation, culture dish was washed 3x with trypsin inside well, rotated 180 degrees, and washed another 3x. Trypsinization was then quenched by adding 160 μL of culture media and total cell mixture was placed into a 1.5mL Eppendorf tube. Cells were spun at 1000gs for 5 minutes, and supernatant was aspirated. Cells were then re-suspended in 200uL of culture media and plated on a 10mm glass bottom dish pre-coated with 25 μg/mL laminin for 1 hr. Cells were then allowed to attach for at least 1 hour, and 2-3 mLs of culture media with or without drug was added to cells.

To trypsinize Axiogenesis hiCMs, manufacturers recommendations were used, as follows. Cells were washed 2x with 500μL PBS*. Cells were placed in 37° C incubator for 7 minutes in PBS*. Following 7 minutes, PBS* was aspirated and 40 μL 0.5% Trypsin (Invitrogen) was placed on cells for 3 minutes in 37° C incubator. Following 3 minutes, 160 μL full Cor.4u media was used to quench trypsinization and re-suspend cells. Cells were then plated on pre-coated glass bottom dish and media added 1.5 hrs later to dilute trypsin and fill chamber. Note* this trypsinization protocol has since been modified by Axiogenesis (now Ncardia). See manufacturer for new protocol.

It is important to note that the trypsinization protocol is based off of the hiCMs manufacturers protocols for cell plating, and these hiCMs have been trypsinized prior to functional assays of cardiomyocyte performance and characterization (e.g., drug response, electro-physiology, maturation, etc…) (Fine et al., 2013; Ivashchenko et al., 2013; Mioulane et al., 2012).

#### Transient Transfection of hiCMs

Cellular Dynamics hiCMs were transfected via modification of manufacturers recommendations as follows. Volumes used are for transfection in 96 well plates. 2 μL of total 200 ng plasmid (containing fluorescently tagged protein of interest, diluted in opti-mem) and 2 μL 1:5 diluted Viafect (Promega, E4981, in opti-mem) was added to 6 μL opti-mem. Entire mixture of 10 μL was added to single 96 well of hiCMs containing freshly exchanged 100 μL full culture media. Transfection was allowed to go overnight (~15 hrs), and washed 2x with full culture media. For transfection of multiple probes, 2 μL of 200ng plasmid was used for each probe together with 4 μL 1:5 diluted Viafect, into 2 μL Opti-mem and mixture was applied to cells as above.

Axiogenesis hiCMs were transfected via modification of manufacturers recommendations as follows. A 3.5:1 Fugene to DNA ratio was used to transfect Axiogenesis hiCMs. 1.2 μL Fugene + 0.33 μgDNA per 96 well into 5μL serum free Cor.4u media was incubated for 15 minutes at room temperature. 95 μL full Cor.4u media was added to mixture and entire mixture was added to hiCMs (on top of 100 μL already in well). For transfection of 2 separate plasmids, 3.5:1 ratio was used for both plasmids and additional volume was subtracted from 95 μL dilution. Note* this transfection protocol has since been modified by Axiogenesis. See manufacturer for new protocol.

#### Protein Knockdown

Cellular Dynamics hiCMs were used for knockdown of NMIIA, NMIIB, and FHOD3 via modification of manufacturers recommendations. Volumes used are for siRNA application in 96 well plates. Dharmacon SmartPool siRNA (GE Dharmacon, Lafayette, CO) targeted to MYH9 (NMIIA), MYH10 (NMIIB), and FHOD3 were used (E-007668-00-0005, E-023017-00-0010, and E-023411-00-0005, respectively). To achieve KD, a master mixture of 100 μl Opti-MEM (ThermoFisher, Waltham, MA) + 4 μl Transkit-TKO (Mirus Bio, Madison WI) + 5.5 μl 10μM siRNA was incubated for 30 min at room temperature. 80 μl of fresh, pre-warmed media was added to hiCMs. Following incubation of siRNA mixture, 8.3 μl of mixture was added to each individual well of 96 well plate. hiCMs were then incubated for 2 days at 37° C. hiCMs were then washed 2x with fresh, pre-warmed media. To achieve KD of NMIIA, NMIIB, and FHOD3, 3 rounds of siRNA mediated KD described above was necessary. Following 3 rounds of treatment, hiCMs still beat and maintained sarcomere structure as shown in Supplemental Video 1 and 2.

**Western Blotting**. Gel samples were prepared by mixing cell lysates with LDS sample buffer (Life Technologies, #NP0007) and Sample Reducing Buffer (Life Technologies, #NP00009) and boiled at 95°C for 5 minutes. Samples were resolved on Bolt 4-12% gradient Bis-Tris gels (Life Technologies, #NW04120BOX). Protein bands were blotted onto a nylon membrane (Millipore). Blots were blocked using 5% NFDM (Research Products International Corp, Mt. Prospect, IL, #33368) in TBST. Antibody incubations were also performed in 5% NFDM in TBST. Blots were developed using the Immobilon Chemiluminescence Kit (Millipore, #WBKLS0500).

#### Fixation and immunohistochemistry

hiCMs and HeLa cells were fixed with 4% paraformaldehyde (PFA) in PBS at room temperature for 20 min and then extracted for 5 min with 1% Triton X-100 and 4% PFA in PBS as previously described (Burnette et al., 2014). Cells were washed three times in 1× PBS. To visualize endogenous βCMII and transfected proteins in fixed cells, hiCMs were live-cell extracted before fixation as described previously to reduce background and non-cytoskeletal myosin II filaments (i.e., the soluble pool). Briefly, a cytoskeleton-stabilizing live-cell extraction buffer was made fresh containing 2 ml of stock solution (500 mM 1,4-piperazinediethanesulfonic acid, 25 mM ethylene glycol tetraacetic acid, 25 mM MgCl2), 4 ml of 10% polyoxyethylene glycol (PEG; 35,000 molecular weight), 4 ml H2O, and 100 μl of Triton X-100, 10 μM paclitaxel, and 10 μM phalloidin. Cells were treated with this extraction buffer for 1 min, followed by a 1-min wash with wash buffer (extraction buffer without PEG or Triton X-100). Cells were then fixed with 4% PFA for 20 min. After fixation, the following labeling procedures were used: for actin visualization, phalloidin-488 in 1 × PBS (15 μl of stock phalloidin per 200 μl of PBS) was used for 3 h at room temperature. For immunofluorescence experiments, cells were blocked in 10% bovine serum albumin (BSA) in PBS. Primary antibodies were diluted in 10% BSA. NMIIA antibody (BioLegend, P909801) was used at 1:1000. NMIIB antibody (Cell Signaling, 3404S and BioLegend 909901) were used at 1:200. βCMII antibody (Iowa Hybridoma Bank, A4.1025) was used undiluted from serum. α-actinin 2 (Sigma-Aldrich, clone EA-53) antibody was used at 1:200. FHOD3 antibody (Santa Cruz Biotechnology, G-5, sc-374601) was used at 1:100. Cardiac Troponin T antibody (Santa Cruz Biotechnology, CT3, sc-20025) was used at 200 μg/ml. Secondary antibodies were diluted in 10% BSA at 1:100 and centrifuged at 13,000 rpm for 10 min before use. Cells were imaged in VECTASHIELD Antifade Mounting Media with DAPI (H-1200, VECTOR LABORATORIES, Burlingame, CA, USA).

#### Structured illumination microscopy

Two SIM imaging modalities were used in this study (individual experiments were always imaged using the same modality). SIM imaging and processing was performed on a GE Healthcare DeltaVision OMX equipped with a 60×/1.42 NA oil objective and sCMOS camera at room temperature. SIM imaging and processing was also performed on a Nikon N-SIM structured illumination platform equipped with an Andor DU-897 EMCCD camera and a SR Apo TIRF (oil) 100x 1.49 NA WD 0.12 objective at room temperature.

#### Spinning Disk Microscopy

Spinning disk confocal images for Figures 1G, 1H, and 3E were taken on a Nikon Spinning Disk equipped with a Yokogawa CSU-X1 spinning disk heard, Andor DU-897 EMCCD camera and 100x Apo TIRF (oil) 1.49 NA WD 0.12mm objective at 37 degrees C and 5%CO_2_.

#### Confocal Microscopy

Confocal images for Figures 1F, 7B, 7C and 7D were taken on a Nikon A1R laser scanning equipped with a 60x/1.40 Plan Apo Oil objective at 37 degrees C and 5%CO_2_. Confocal images for Fig. 3e and Supplemental Fig. 6d were taken on a Zeiss 880 with AiryScan equipped with a 63x/1.40 Plan-Apochromat Oil objective at room temperature.

#### Wide-field Microscopy

Images for data which led to Figure 1D were taken on a high-resolution wide-field Nikon Eclipse Ti equipped with a Nikon 100x Plan Apo 1.45 NA oil objective and a Nikon DS-Qi2 CMOS camera. Videos 1 and 2 were taken on a Leica DMIL cell culture microscope equipped with a N PLAN L 40x/0.55 CORR PH2 at room temperature.

#### Antisense morpholino oligonucleotide injections

For zebrafish morpholino knockdown experiments, splice-blocking antisense morpholinos (MO) oligonucleotides were injected into singe-cell wild-type embryos. Embryos were allowed to develop until 48 hours post fertilization (hpf) and then fixed for analysis.

Control experiments have been previously published to confirm morpholino efficacy and specificity including; examination of the morpholino effect on the *myh10* transcript using RT-PCR, rescue of the *myh10* morpholino knockdown phenotype with expression of human *myh10* mRNA, and Western blot analysis of morpholino effects on NMIIB protein levels (Gutzman et al., 2015).

Morpholino details are as follows:

*myh10* MO (5’–CTTCACAAATGTGGTCT-TACCTTGA-3’; Gene Tools) targets EXON2-intron2.

Standard control MO (5’–CCTCTTACCTCAGTTACAATTTATA-3’; Gene Tools). Zebrafish *p53* MO (5’ –GCGCCATTGCTTTTGCAA-GAATTG-3’; Gene Tools). p53 MO was used in conjunction with either standard control MO or *myh10* MO. 3ng of each morpholino was used for all experiments. 25 pg of mCherry mRNA was co-injected with each condition to screen for injection efficiency.

### QUANTIFICATION AND STATISTICAL ANALYSIS

#### Data Quantification

To quantify % of hiCMs with sarcomeres (i.e., Figure 2B), the actin cytoskeleton (via fluorescently labeled phalloidin) was imaged using structured illumination microscopy. hiCMs were quantified as containing sarcomere structures if they contained at least 1 myofibril containing at least 3 Z-discs (bright phalloidin staining which overlaps with α-actinin 2 staining as in Figure 2B) spaced ~1.5-2 μm apart. By these metrics our quantification of sarcomere formation is not a measure of sarcomere maturity or alignment, but a measure of the hiCMs ability to assemble the building blocks of the sarcomere (i.e., the thin actin filaments) in response to a perturbation. Thus, while NMIIA KD hiCMs clearly form unaligned, disorganized, and fewer sarcomeres and myofibrils than control hiCMs, they still maintain the ability to assemble sarcomere structures, which is reflected in our quantification (Figures 4E and 4G). In the same vein, we realize our quantification is a very liberal quantification of sarcomere assembly. While we don’t expect a small array of sarcomeres to represent a functionally capable cardiomyocyte, we are investigating the early steps of sarcomere assembly and the ability of hiCMs to form the basic actin-myosin structure of the sarcomere. A similar methodology was used to quantify βCMII A-band filament stacks in Fig. 3g-h, using endogenous βCMII staining and SIM instead of the actin cytoskeleton. hiCMs were quantified as containing βCMII A-band filament stacks if they contained even one βCMII filament stack, comparable to the smallest βCMII filament stacks found in control hiCMs. Sarcomere formation in Zebrafish experiments (Fig. 2h) is described above. Briefly, the bright actin staining representing Z-discs was used to count all Z-discs present in control and *myh10* MO animals. The number of Z-discs per image were divided by the size in microns of that individual animal heart for the Z-disc/μm measurement in Fig. 2h. For MSF retrograde flow rates (Fig. 1 g-i), 3 regions of interest (ROIs) were used per cell. ROIs were drawn using the line tool in FIJI starting from in front of the leading edge (to ensure new MSF formation was captured) to the cell body where sarcomere structures were localized. ROIs were then used to measure MSF translocation rates using the kymograph tool (line width = 3) on hiCMs which had been aligned using StackReg function. Kymographs generated in this manner were then manually measured by counting pixels on the X axis (distance) and the Y axis (time) for a distance/time measurement resulting in translocation rates. This method is similar to previously described methods of actin arc translocation in non-muscle cells.

#### Actin staining and sarcomere quantification

48 hours post fertilization (hpf), embryos were fixed in 4% paraformaldehyde in PBT for 2 hours or overnight at 4°C. Embryos were then washed three times in PBT. To visualize actin, embryos were incubated in Alexa Fluor 488 phalloidin (Invitrogen A12379) at a ratio of 1:40 in PBT at 4°C for 2 days. Next, embryos were washed five times in PBT. For imaging, embryos were manually deyolked and heart tissue dissected. Heart tissue was mounted on a microscope slide in 100% glycerol for imaging. Imaging was conducted using a Nikon CS2 laser-scanning confocal microscope with NIS Elements software. Maximum projections were created by the last author of each ventricle and each image file was labeled by a random number. The first author then marked each sarcomere in a blinded fashion.

#### Immunofluorescence localization quantification

To quantify localization of NMIIA, NMIIB, and βCMII, line scans starting from the edge of the cell were taken for every cell and the normalized average localizations were used to average the number of indicated experiments for the final localization patterns depicted in the graph (as in Figure 4B).

To quantify Arp2/3 intensity, a similar but altered strategy was taken. 4 separate boxes for each cell were placed along the edge of the cell (i.e., the lamellipodium) using the actin channel for guidance. These boxes were then used to measure average intensity of the anti-p34 channel (i.e., the Arp2/3 complex). Background subtracted averages for each cell in control and CK666 treated hiCMs were used to quantify percent decrease as depicted in Figure 2D.

#### Cells Used for this Study

In order to have comparable results, cells used for this study were standardized based on morphology. Specifically, spread non-muscle cells and hiCMs with a broad leading edge and lamella were chosen as previously shown in studies of both non-muscle and muscle contractile system formation (Burnette et al., 2014; Rhee et al., 1994). This also facilitated the ability to observe the MSF to sarcomere transition in live hiCMs, and is recommended for future studies investigating sarcomere assembly. All experiments measuring sarcomere assembly were conducted at least 3 times, separately, and cells were imaged using SIM. Though this results in a relatively small number of cells for some of the experiments, we believe super-resolution imaging modalities such as SIM offer invaluable insight into sarcomere assembly (Gustafsson, 2005). Indeed, sub-diffraction imaging is required to reliably localize myosin II co-filaments both *in vitro* and *in vivo*. Furthermore, as has been seen in *Drosophila,* even high-resolution imaging modalities such as laser-scanning confocal microscopy are not sufficient to detect subtle, yet important, structural changes in response to perturbation (Fernandes and Schock, 2014).

#### Statistics

Statistical significance was calculated using 2-tailed, unpaired Students T-tests performed in Excel. Error bars in all graphs represent standard error of the mean (SEM). For all graphs depicting % of cells (for example, Figure 1D), number of cells and experiments is indicated in figure legend. Percents displayed represent the average of the averages of the all experiments performed. For example, if controls cells displayed 95%, 90%, and 100% of all cells displaying actin arcs in 3 separate experiments, the represented percentage in the graph would be 95%. SEM was calculated by dividing the standard deviated by the square root of the number of separate experiments. For actin translocation rates at least 3 measurements per cell were used to calculate the average translocation rates per cell. Translocation rates were then calculated in same manner as described above. Line-scan graphs represent normalized relative fluorescence.

## Author Contributions

D.T.B conceived the study. A.M.F. and D.T.B designed the research. A.F.M conducted a majority of experiments. D.T.B, A.M.F, A.C.N., and N.T. analyzed the data; N.T. and A.C.N. assisted in hiCM and non-muscle cell experiments, M.R.V. and J.H.G. performed Zebrafish MO and imaging experiments, and S.W.C. and M.J.T. assisted in creating βCMII-mEGFP construct. A.E.M. and D.M.B. acquired mouse tissue samples. B.R.N. and J.R.B. acquired human tissue samples. A.M.F. and D.T.B. wrote the manuscript with input from all authors.

## Acknowledgements

We thank the M. Tyska lab and the Epithelial Biology Center (EBC) at Vanderbilt University for invaluable discussions and reagents. Janice Williams of the Cell Imaging Shared Resources was instrumental in acquiring the EM images in Figure 1A. We give a special thanks to Sean Schaffer and Bryan Millis of the Cell Imaging Shared Resources and Nikon Center of Excellence at Vanderbilt University for help with and maintenance of the SIM and spinning disk microscopes, and Anthony Tharp and Josh Luffman for the CDB CORE Equipment. The authors declare no competing financial interests.

This work was supported by Vanderbilt University School of Medicine Molecular Biophysics Training Grant T32 GM08320 to A.M.F, an American Heart Association pre-doctoral fellowship #16PRE29100014 to A.M.F., a NIH F31 pre-doctoral fellowship # 1 F31 HL136081-01 to A.M.F., a Career Development Award from the National Cancer Institute SPORE in GI Cancer P50 CA095103 to D.T.B., an American Heart Association Scientist Development Grant #17SDG33460353 to D.T.B., and a MIRA Grant (R35) GM125028-01 from NIGMS to D.T.B.

## Supplemental Movie Legends

**Supplemental Movie 1: Control siRNA treated hiCM assembling sarcomeres following plating**

hiCM treated with control siRNA and transfected with Lifeact-mApple and imaged starting at 3 hours post plating. Note increase in actin signal as movie proceeds. MSFs undergo retrograde flow and transition to sarcomere containing myofibrils towards cell body as in untreated control hiCMs (i.e., control cells not treated with Non-targeting siRNA). 46.87 by 46.87 μms. Video length 14.5 hours, 5fps.

**Supplemental Movie 2: NMIIB KD hiCM displays prominent actin dynamics defects**

hiCM treated with siRNA *myh10* (NMIIB KD), transfected with Lifeact-mApple and imaged starting at 3 hours post plating. Note almost complete absence of actin retrograde flow and excessive protrusion and retraction events at edges of NMIIB KD hiCM compared with control hiCM (Supplemental Movie 1). 46.87 by 46.87 μms. Video Length 16.5 hours, 5 fps.

**Supplemental Movie 3: siRNA Scramble hiCMs beating in 96 well plate pre-trypsinization**

hiCMs 3 days post treatment with siRNA scramble. hiCMs beat comparably to untreated control cells. Phase contrast, 20x.

**Supplemental Movie 4: siRNA MYH10 (NMIIB KD) hiCMs beating in 96 well plate pre-trypsinization**

hiCMs 3 days pot treatment with siRNA MYH 10 (for NMIIB KD). hiCMs beat comparably to siRNA scramble control hiCMs as shown in Video 3. Phase contrast, 20x.

**Figure S1:**
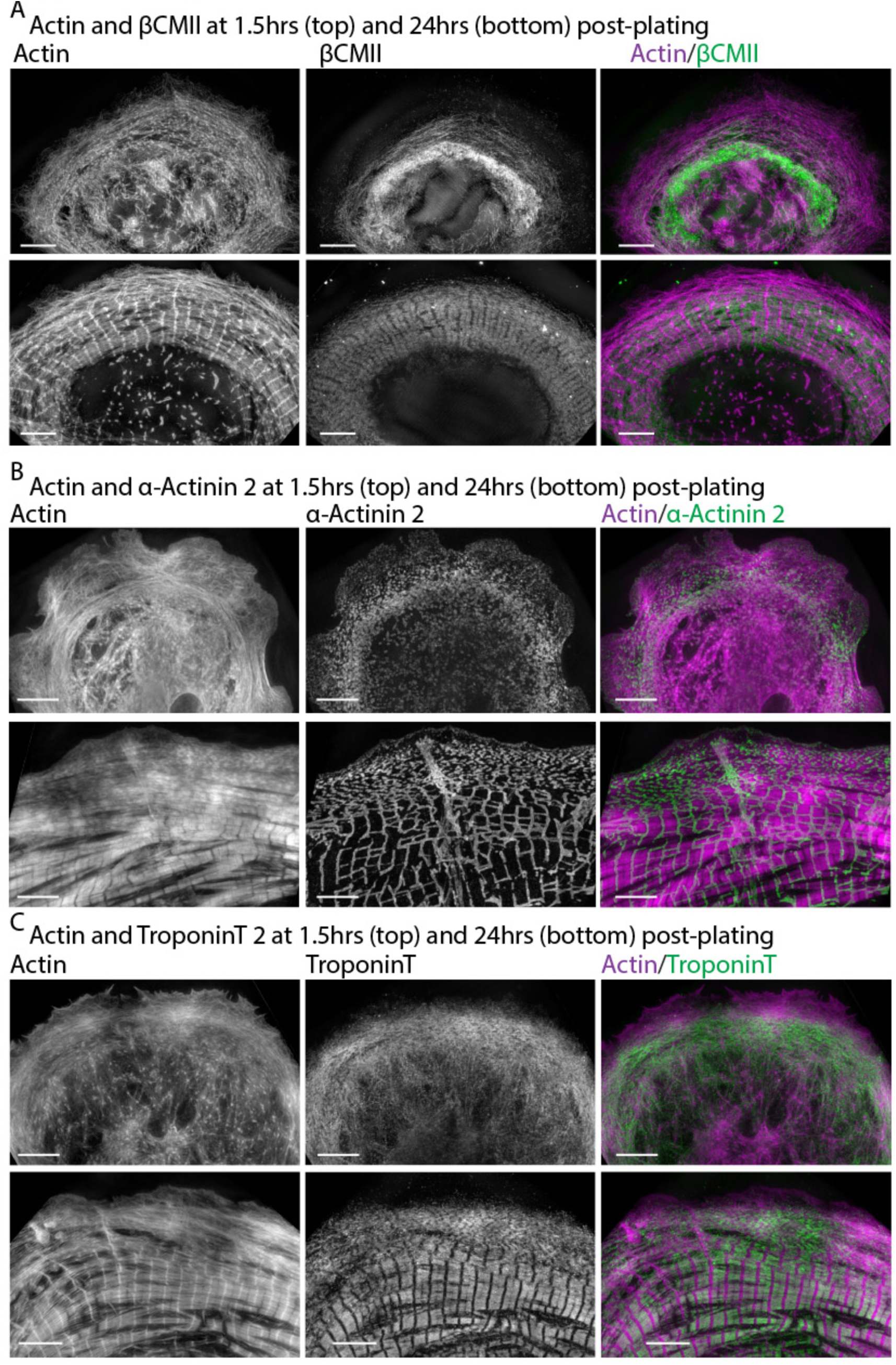
hiCMs don’t contain sarcomeres at early time points post plating. (A) hiCM stained for actin and βCMII at 1.5hrs (top) and 24 hrs (bottom) following trypsinization. (B) hiCM stained for actin and α-actinin 2 at 1.5 hrs (top) and 24hrs (bottom) following trypsinization. (C) hiCM stained for actin and TroponinT at 1.5 hrs (top) and 24hrs (bottom) following trypsinization. Note how hiCMs at both 1.5 hrs contain Muscle Stress Fibers (MSFs), but contain neither actin nor βCMII based sarcomeres. hiCMs at 24hrs however contained prominent sarcomere structures. Scale bars, 5 μm.

**Figure S2.**
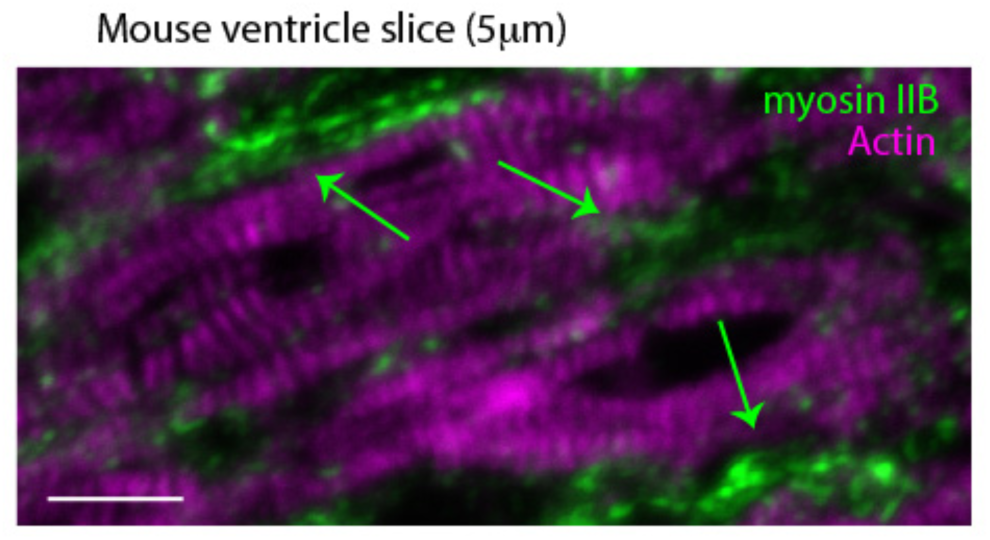
NMIIB localization *in vivo*. Localization of NMIIB and actin in tissue slice from heart ventricle of P3 mouse. Green arrows denote strong localization of NMIIB outside of but adjacent to sarcomere structures as in hiCMs (Figure 4A). Scale Bar: 10 μm.

**Figure S3:**
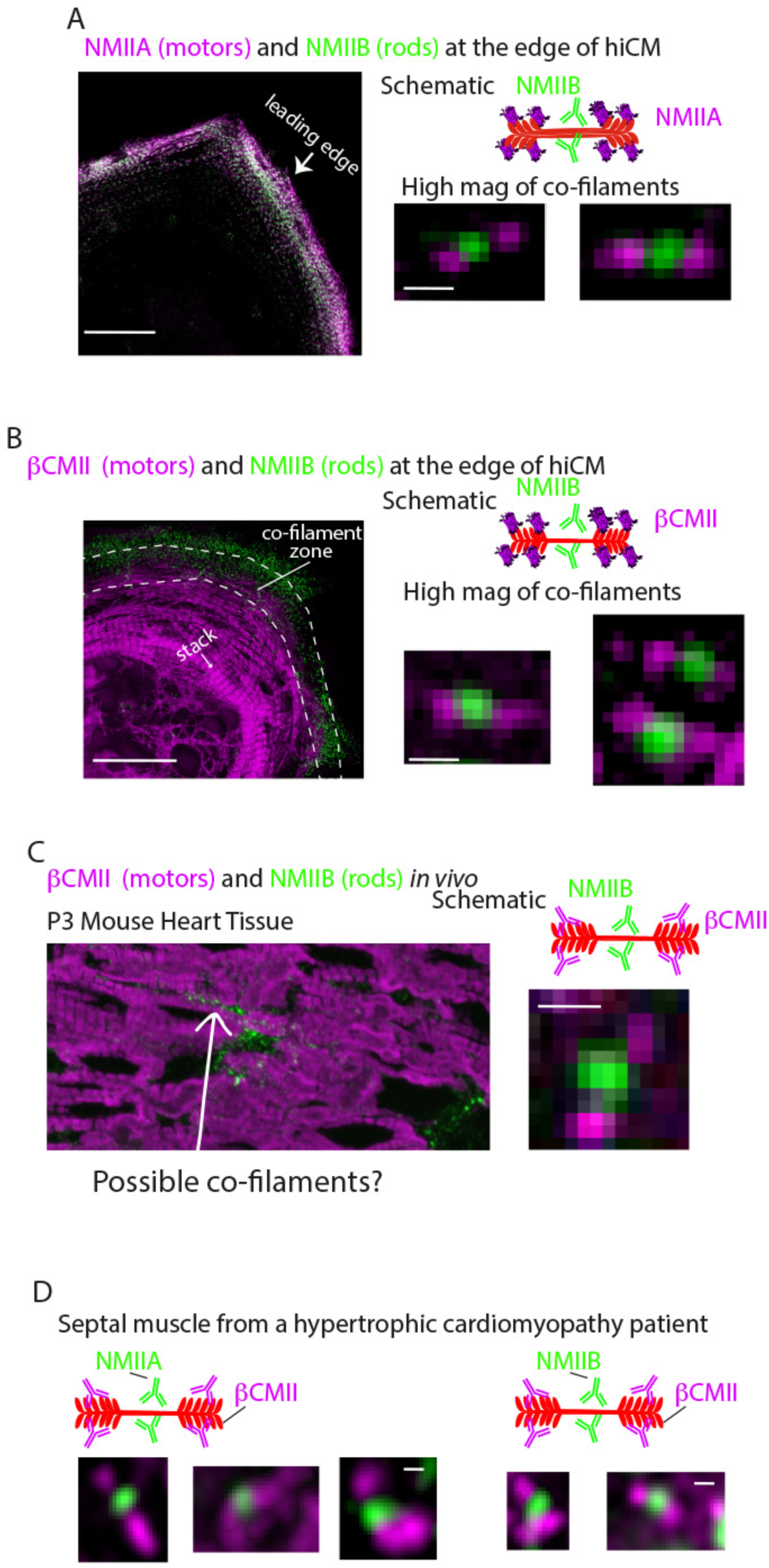
Myosin II co-filaments *in vitro* and *in vivo*. (A) hiCM from Figure 4C transfected with NMIIA-mEmerald (N-terminal motors) and stained for NMIIB rods. Schematic depicts visualization strategy. Low mag of hiCM left, and high mag examples bottom right. (B) hiCM transfected with βCMII-mEGFP (N-terminal motors) and stained for endogenous NMIIB rods. Schematic depicts visualization strategy. Low mag of hiCM left, and high mag examples bottom right. (C) Low mag view of P3 mouse heart tissue stained for βCMII (motors) and NMIIB (rods). As in hiCMs, NMIIB is restricted from sarcomere structures but is localized adjacently to sarcomeres. Arrow indicates area of possible co-filaments. High mag example shown at right is taken from P3 mouse tissue imaged on Zeiss 880 with AiryScan from similar area indicated by arrow on low mag image. Schematic indicates visualization strategy. (D) High mag examples of NMIIA (left) and NMIIB-βCMII co-filaments from human hypertrophic cardiomyopathy patients imaged on Zeiss 880 with AiryScan. Cartoon schematic indicates imaging strategy. Scale bars: (A), (B), 10 μm (left); (A), (B), (C), (D), 200 nm (right).

**Supplementary Figure S4:**
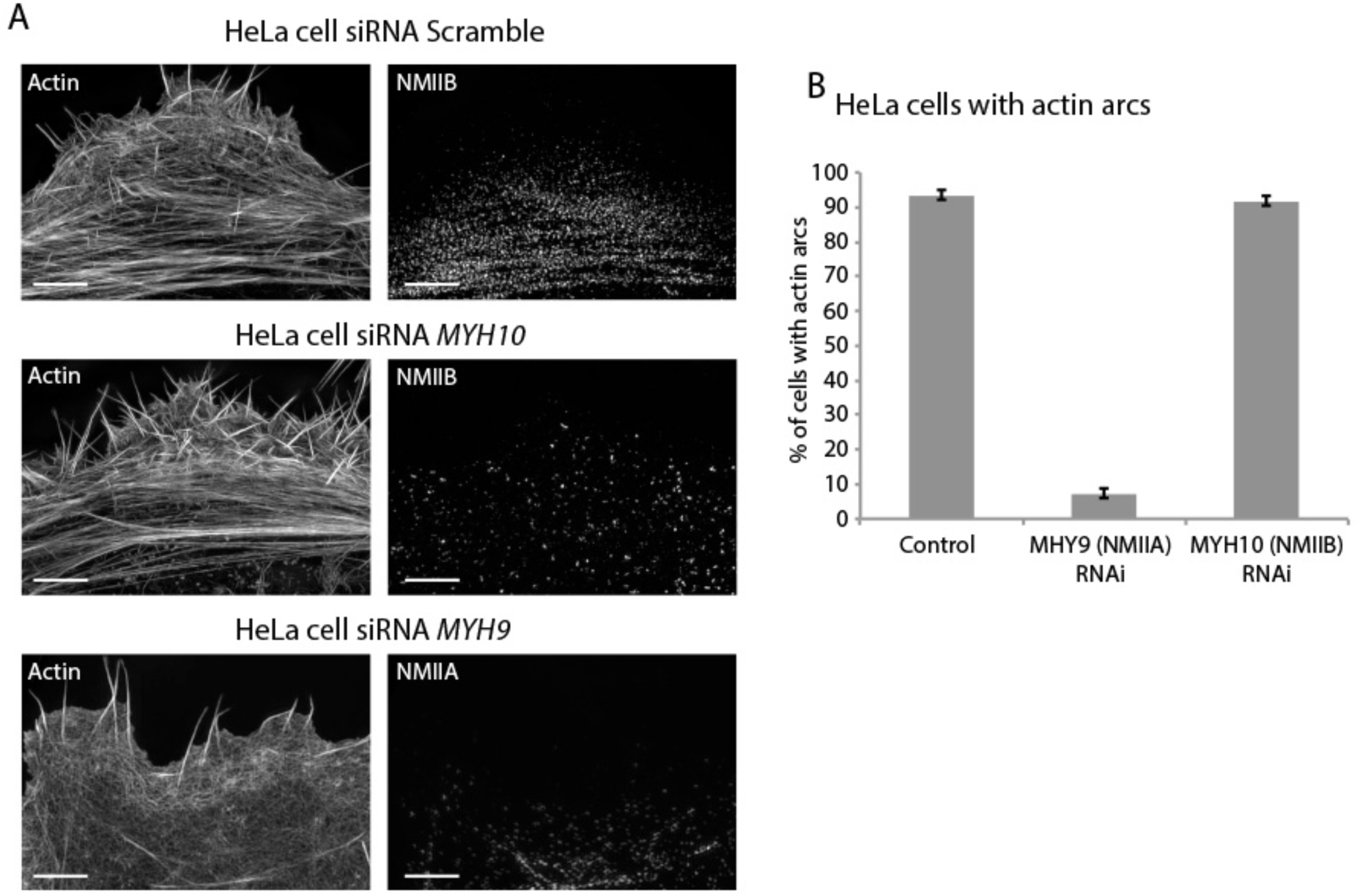
NMIIA but not NMIIB is required for actin arc formation in HeLa cells. (A) HeLa cells treated with control, anti *myh10*, or anti *myh9* (top, middle, bottom, respectively) siRNA. Note how control and NMIIB KD cells display prominent actin arc stress fibers, but NMIIA KD cells contain no actin arcs. (B) Quantification of HeLa cells with actin arc stress fibers in indicated conditions. Scale Bars: 5 μms.

**Figure S5:**
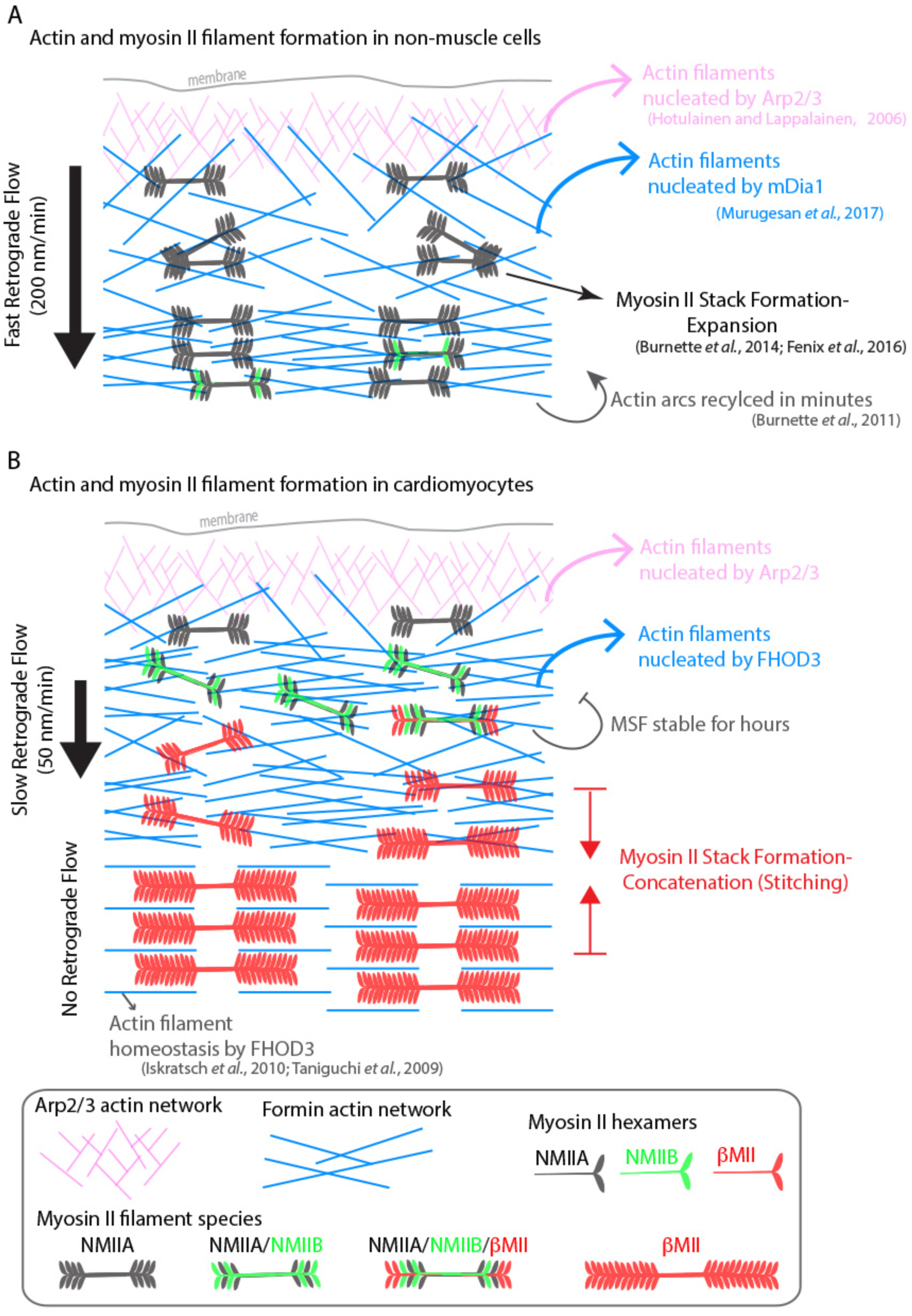
Model of actomyosin stress fiber formation in non-muscle cells and human cardiomyocytes. Actin and myosin II stress fiber formation in non-muscle cells. Actin stress fibers are formed via the Arp2/3 complex and the formin mDia1. NMIIA is the predominant isoform at the leading edge of non-muscle cells, and stress fiber formation is NMIIA dependent. Non-muscle cells display robust retrograde flow of actin stress fibers and display rapid turnover. Large NMIIA stacks are formed via growth and expansion of smaller NMIIA filaments. Citations leading to this model are presented in the cartoon. (B) Model of actin and myosin II stress fiber formation in human cardiomyocytes. Sarcomeres are templated by Muscle Stress Fibers (MSFs). MSFs do not require the Arp2/3 complex, and require the formin FHOD3. MSFs display slow retrograde flow compared with non-muscle stress fibers. Both NMIIA and NMIIB are localized to the edge of hiCMs, and display prominent NMII co-filaments. NMIIB-βCMII co-filaments are also present with MSFs. Large βCMII filament stacks form via concatenation and stitching of individual βCMII filaments.

